# Gut microbiota of ring-tailed lemurs (*Lemur catta*) vary across natural and captive populations and correlate with environmental microbiota

**DOI:** 10.1101/2021.06.27.450077

**Authors:** Sally L. Bornbusch, Lydia K. Greene, Sylvia Rahobilalaina, Samantha Calkins, Ryan S. Rothman, Tara A. Clarke, Marni LaFleur, Christine M. Drea

## Abstract

**Background:** Inter-population variation in host-associated microbiota reflects differences in the hosts’ environments, but this characterization is typically based on studies comparing few populations. The diversity of natural habitats and captivity conditions occupied by any given host species has not been captured in these comparisons. Moreover, intraspecific variation in gut microbiota, generally attributed to diet, may also stem from differential acquisition of environmental microbes – an understudied mechanism by which host microbiomes are directly shaped by environmental microbes. To more comprehensively characterize gut microbiota in an ecologically flexible host, the ring-tailed lemur (*Lemur catta*; n = 209), while also investigating the role of environmental acquisition, we used 16S rRNA sequencing of lemur gut and soil microbiota sampled from up to 13 settings, eight in the wilderness of Madagascar and five in captivity in Madagascar or the U.S. Based on matched fecal and soil samples, we used source-sink ecological theory to examine covariation between the two types of consortia.

**Results:** The diversity of lemur gut microbes varied markedly within and between settings. Microbial diversity was not consistently greater in wild than in captive lemurs, indicating that this metric is not necessarily an indicator of host habitat or condition. Variation in microbial composition was inconsistent with either a single, representative gut community for wild conspecifics or a universal ‘signal of captivity’ that homogenizes the gut consortia of captive animals. Despite the similar, commercial diets of captive lemurs on both continents, lemurs within Madagascar were compositionally most similar, suggesting that non-dietary factors govern some of the variability. In particular, soil microbial communities were most distinct between the two continents, and there was significant and context-specific covariation between lemur gut and soil microbiota.

**Conclusions:** As one of the broadest, single-species investigations of primate microbiota, our study highlights that gut consortia are sensitive to multiple scales of environmental differences. This finding begs a reevaluation of the simple ‘captive vs. wild’ dichotomy. Beyond the important implications for animal care, health, and conservation, our finding that environmental acquisition may mediate aspects of host-associated consortia further expands the framework for how host-associated and environmental microbes interact across different microbial landscapes.

## Introduction

The structure of gut microbial communities within vertebrates is influenced in part by endogenous host factors, such as genotype and physiology^1–3^, and in part by exogenous factors, such as sociality, seasonality, habitat quality, and diet^4–6^. These exogenous factors can influence which microbial taxa in a gut community thrive or become depauperate, as amply demonstrated in dietary studies^7–10^, or they can provide opportunities for more direct routes of microbial acquisition^11–14^. For example, the transmission of microbes between hosts, including horizontal pathogen transfer^15–17^ or vertical transmission during the birthing process and nursing^18,19^, are significant drivers of host health and development. There is, likewise, the potential for horizontal acquisition of microbes via exposure to environmental consortia on natural (e.g., soil) and man-made surfaces, plus on food and in water^12,20–23^; however, this latter route to shaping host-associated communities, hereafter referred to as ‘environmental acquisition,’ remains understudied. Here, we match-sampled ring-tailed lemur (*Lemur catta*) feces and soil from 13 settings, representing both a large portion of the lemurs’ natural habitat range in Madagascar and a range of captive housing conditions in Madagascar and the U.S. (Table 1), to (a) characterize variation in host gut microbiota, (b) characterize variation in soil microbiota, and (c) test for any covariation between host and soil communities. Examining environmental microbes alongside host-associated communities is a first step to understanding the role of environmental acquisition in population-level differences between host microbiomes.

**Table 1.** Research settings (names, descriptions, and locations) and samples collected under wilderness conditions and under captivity conditions in Madagascar and the U.S. A subset of the samples collected were omitted from analyses owing to low-yield extractions or low-quality sequencing. Soil samples could not always be obtained. Maps show locations of each setting; the gray shaded area shows the natural range of wild ring-tailed lemurs in Madagascar.

| Condition | Setting (abbreviation) | Setting description | Samples:<br>collected (analyzed) |  |
| --- | --- | --- | --- | --- |
|  |  |  | fecal | soil |
| wilderness | Amoron'I Onilahy (AMO) | Riverine gallery forest, dry scrub forest | 20 | 4 (3) |
|  | Berenty Reserve (BER) | Semi-arid dry forest, spiny forest | 22 (19) | 4 |
|  | Beza Mahafaly Special Reserve (BEZ) | Riverine gallery and semi-arid spiny forest | 26 | 4 (3) |
|  | Fiheranana (FIH) | Dry forest and spiny forest | 2 | - |
|  | Isalo National Park (ISO) | Dry deciduous forest | 18 (16) | 3 |
|  | Ivohiboro (IVO) | Humid forest, grassland | 15 | - |
|  | Ranomay (RAN) | Dry forest | 13 | 2 (1) |
|  | Tsimanampetsotsa National Park (TSI) | Dry forest and spiny forest | 29 (28) | 8 |
| captivity:<br>Madagascar | Lemur Rescue Center (Toliara, Madagascar; LRC) | Outdoor enclosures in dry and spiny forest | 33 | 2 (1) |
|  | Various towns (pets) | Pets housed in human dwellings | 8 | - |
| captivity:<br>U.S. | Duke Lemur Center (Durham, NC; DLC) | Indoor and outdoor enclosures, including free ranging in semi-deciduous forest | 22 | 3 (2) |
|  | National Zoological Park (Washington, DC; NZP) | Indoor and outdoor enclosures, moated island with vegetation | 4 | - |
|  | North Carolina Zoo (Asheboro, NC; NCZ) | Indoor and outdoor enclosures, moated island with vegetation | 3 | - |
| Total samples: |  |  | 215 (209) | 30 (25) |

Madagascar
wilderness:
- BEZ - FIH - AMO - BER - RAN - TSI - IVO - ISA
captivity: Madagascar
- pet - LRC
captivity: U.S.
- NCZ - DLC - NZP

Previous studies of intraspecific variation in gut microbiota, often framed using a ‘wild vs. captive’ comparison, have provided valuable descriptions of differences in presumed extremes^24–26^. For example, researchers often report a ‘signal of captivity,’ whereby the gut microbiota of captive hosts differ significantly from those of wild conspecifics, converging on a perturbed or ‘humanized’ composition^25,27,28^. Perturbations of this nature are generally attributed to commercial diets that include manufactured chow and cultivated produce^27,29,30^; nevertheless, studies of captive populations have been focused on accredited zoos or rescue facilities that may not represent the range of captive conditions or may be confounded by within-species comparisons across continents^26,29,31^. Even comparative field studies have been limited in the number of populations per species studied, typically to a few populations that differ on a given metric of interest (e.g. season, habitat type or quality^32–34)^. Because hosts experience a wider range of environmental settings than is typically encompassed within wild vs. captive comparisons, a broader comparative approach is necessary to provide a more comprehensive and nuanced understanding of gut microbiota variation.

As noted, differential exposure to environmental microbes provides potential for horizontal transmission and environmental acquisition^20,22,23, 35–37^, with the ingestion of specific microbes being linked to novel digestive functions of the gut microbiota^38–40^. Under certain conditions, environmental acquisition has been shown to outweigh vertical transmission as the main mode of microbial colonization^41,42^. Although environmental acquisition may promote heterogeneity within and between hosts^43^, its role rarely has been considered a differentiating factor between wild and captive hosts. Husbandry practices and veterinary care, for example, introduce cleaning products and antibiotics to the microbial environment of captive animals^44,45^, further differentiating it from natural habitats^46^, with potentially critical consequences to microbiome structure and function.

Ring-tailed lemurs are semi-terrestrial, omnivorous strepsirrhine primates^47,48^ that occupy various habitats across southern Madagascar^49^ and also survive well in captivity^50^. Their ecological flexibility, coupled with existing knowledge about their gut microbiome^26,51–53^, motivates broader comparative study of intraspecific variation that takes environmental acquisition of microbes into consideration. Here, we combine 16S rRNA sequencing and statistical tools based on source-sink ecological theory^54^ to analyze covariation between lemur gut microbiota (e.g., sink communities) and soil microbiota (e.g., source communities). Given the variability of environmental factors across our multiple settings, we expect the diversity, membership, and composition of lemur gut microbiota and soil microbiota to differ within and between three, broad ‘environmental’ conditions: wilderness, captivity in Madagascar, and captivity in the U.S (Table 1).

If diet or habitat quality were the main driver of gut microbiota composition, we would expect (a) wild lemurs to show variation between their natural settings, (b) captive lemurs, regardless of continent, to show similar gut microbiota between their settings (reflecting commercial diets and perturbed habitats), and (c) wild and captive lemurs to differ most drastically from one another, in line with prior studies^27^. If, however, environmental acquisition were to play a major role in shaping lemur gut microbiota, we would again expect (a) wild lemurs to show variation between their natural settings (reflecting the soil microbiota of the lemurs’ habitat), but we would expect (b) Malagasy lemurs (wild and captive) to share certain soil-derived microbiota, differing most drastically from lemurs in captivity in the U.S., and (c) differential access to soil within captive conditions to correlate with differential soil-associated microbes present in hosts. With regard to the latter, for example, we might expect greater proportions of soil-associated microbes in captive lemurs that gain access to natural enclosures compared to their counterparts that are housed indoors.

## Results

### Lemur gut microbiota: Variation in diversity, membership, and composition

#### Alpha diversity

Across the gut microbiota of all ring-tailed lemurs sampled in this study, metrics of alpha diversity differed significantly by environmental condition (Generalized Linear Models or GLM; Shannon: F = 23.773, p < 0.001; Faith’s phylogenetic: F = 4.415, p = 0.013; Figures 1a, b) and by setting (GLM; Shannon: F = 13.157, p < 0.001; Faith’s phylogenetic: F = 5.628, p < 0.001; Figures 1c, d; Supplementary Material 1). The gut microbiota of wild lemurs and captive lemurs in the U.S. (hereafter, ‘captive_U.S._ lemurs’) were similarly diverse overall (pairwise Wilcoxon test; Shannon: p = 0.635; Faith’s phylogenetic: p = 0.056; Figures 1a, b), whereas those of captive lemurs in Madagascar (hereafter, ‘captive_M_ lemurs’) were significantly less diverse (pairwise Wilcoxon test; Shannon, wild vs. captive_M_ lemurs: p < 0.001; wild vs. captive_U.S._ lemurs: p < 0.001; Faith’s phylogenetic, wild vs. captive_M_ lemurs: p = 0.022; wild vs. captive_U.S._ lemurs: p = 0.021; Figures 1a, b). Within environmental condition, however, both metrics of alpha diversity varied widely between the different settings (Figures 1c, d; Supplementary Material 1). For example, among wild lemurs, setting was a significant predictor of both metrics of alpha diversity (GLM; Shannon diversity: F = 20.768, p < 0.001; Faith’s phylogenetic: F = 11.104, p < 0.001). Sex was not a significant predictor in any models of either alpha diversity metric (Supplementary Material 1).

**Figure 1.**
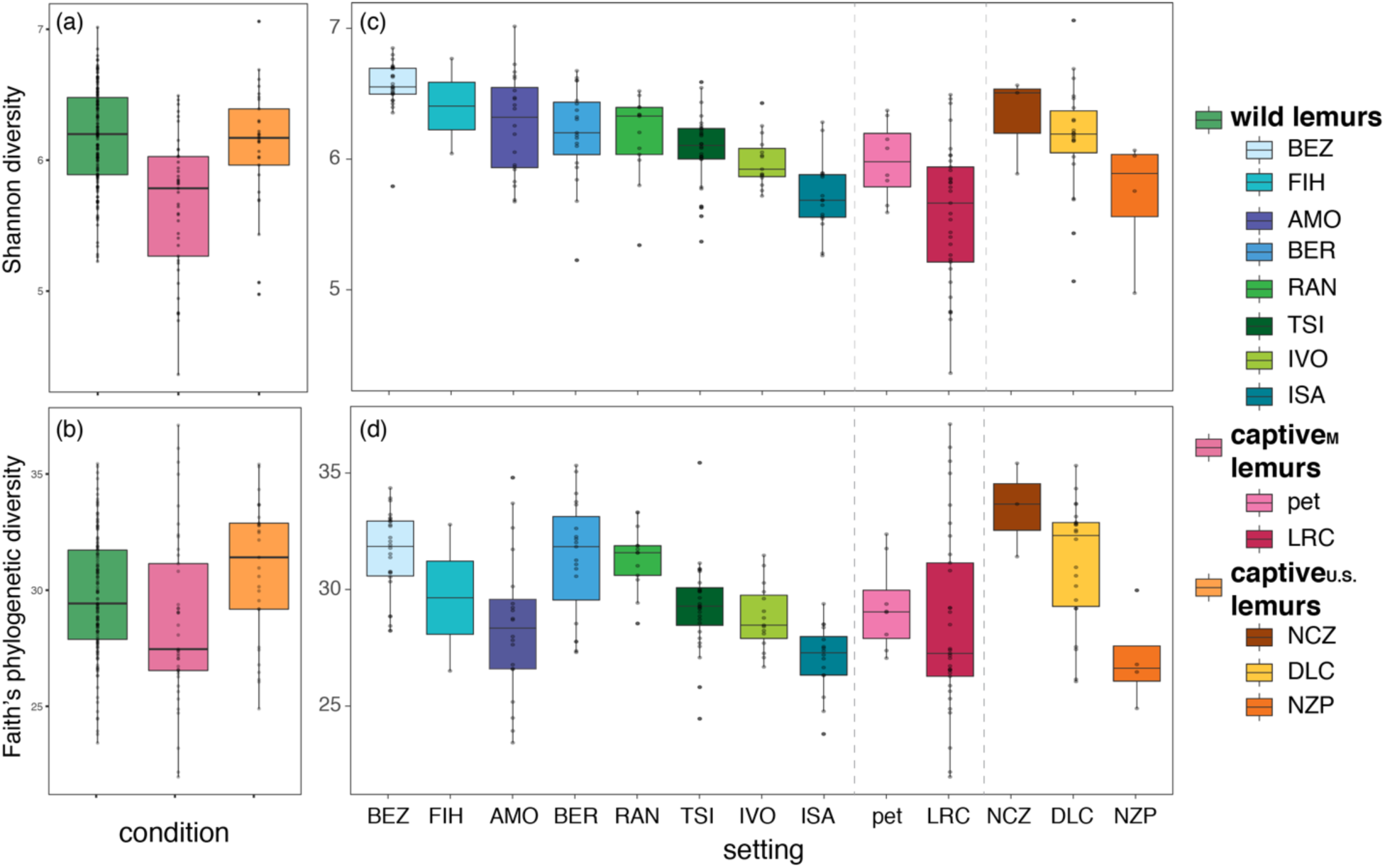
Alpha diversity metrics of lemur gut microbiota (a, b) collapsed by environmental condition, including wilderness (wild lemurs; green), captivity in Madagascar (captive_M_ lemurs; pink), and captivity in the U.S. (captive_U.S._ lemurs; orange), and (c, d) averaged across individuals for each of the 13 different settings inhabited (reprising the color codes of each condition, delineated by dashed vertical lines). Shown are both (a, c) Shannon diversity and (b, d) Faith’s phylogenetic diversity. Across the (c, d) settings within a condition (see Table 1 for names of abbreviated study sites), the data are plotted in descending order of mean Shannon diversity.

#### Community membership

The membership of lemur gut microbiota included 64 abundant taxa (i.e., those that accounted for >1% of sequences). Of these 64 taxa, only four (6.2%) were shared across lemurs from all settings: the genera *Bacteroides* (phylum Bacteroidetes), *Rikenellaceae RC9 gut group* (Bacteroidetes), *Erysipelotrichaceae* UCG-004 (Firmicutes), and *Treponema 2* (Spirochaetes). Within condition, five (7.8%) taxa were shared by all wild lemurs, whereas 10 (15.6%) and six (9.4%) taxa were shared by captive_M_ and captive_U.S._ lemurs, respectively (Figure 2). Using Analysis of Compositions of Microbiomes (ANCOM), we identified 801 amplicon sequence variants (ASVs) that were differentially abundant across the three conditions. For example, members of the Erysipelotrichaceae family characterized the microbiota of wild lemurs, whereas taxa from the Spirochaetaceae and Prevotellaceae families were more abundant in the gut microbiota of captive lemurs from both continents.

**Figure 2.**
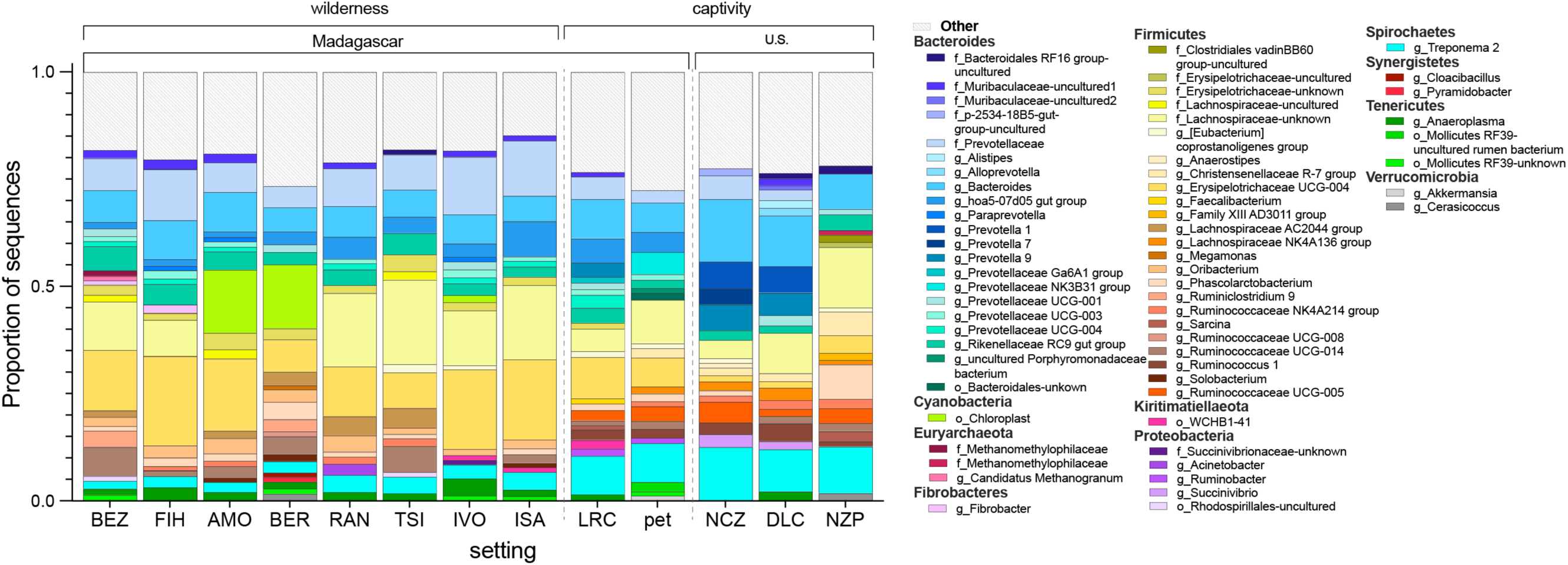
Mean proportion of sequences assigned to microbial taxa across lemurs at each of the 13 different settings, with the three conditions (wilderness, captivity in Madagascar, and captivity in the U.S.) delineated by dashed vertical lines (see Table 1 for names of abbreviated study sites). Taxa are identified by phylum and deepest possible taxonomic level (i.e., genus level or above); those representing < 1% of the microbiomes were combined into the category “Other”

*Erysipelotrichaceae UCG-004* and *Treponema 2*, for example, were abundant in all lemurs (Figure 2), but the log ratios of the two genera distinguished lemur gut microbiota by the three environmental conditions and, in particular, differentiated wild lemurs from captive_U.S._ lemurs (Figure 3).

**Figure 3.**
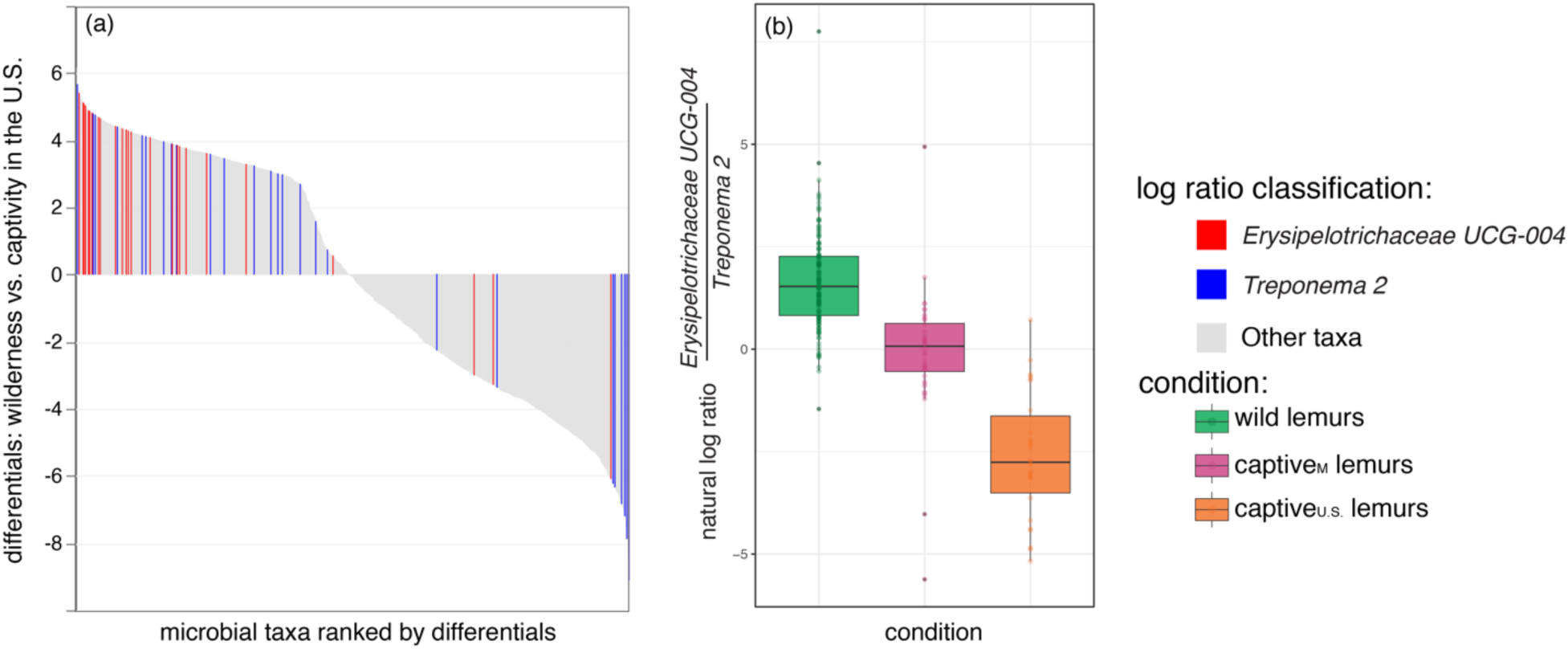
Differential abundance of *Erysipelotrichaceae UCG-004* and *Treponema* 2 in the gut microbiota of lemurs. (a) Differential rank plot showing lemur gut microbial taxa (x axis) ranked by their differentials (y axis; the estimated log-fold changes for taxa abundances across sample groups) for lemurs in the wilderness vs. captivity settings in the U.S. Those taxa that are more abundant in the wild lemurs compared to captive lemurs in the U.S. appear on the right side of the plot whereas those that are less abundant in wild lemurs appear on the left side. *Erysipelotrichaceae UCG-004* and *Treponema 2* differentials are highlighted in red and blue, respectively, with other taxa represented in gray. (b) Natural log ratios of *Erysipelotrichaceae UCG-004* vs. *Treponema 2* in lemurs across all three environmental conditions.

#### Beta diversity

The composition of lemur gut microbial communities was significantly distinct across the three environmental conditions, as revealed by beta diversity (Permutational Multivariate Analysis of Variance or PERMANOVA; wild vs. captive_M_ lemurs: pseudo-F = 30.169, p < 0.001; wild vs. captive_U.S._ lemurs: pseudo-F = 97.912, p < 0.001; captive_M_ vs. captive_U.S._ lemurs: pseudo-F = 20.808, p < 0.001). Across all subjects, gut microbiota composition clustered distinctly by condition (principal coordinate analysis of unweighted UniFrac distances; Figures 4a, b). One notable exception, however, owed to a single pet lemur: Unlike its in-country peers (i.e., other captive_M_ lemurs), its microbial community structure matched those of wild lemurs (see arrows in Figures 4a, b).

**Figure 4.**
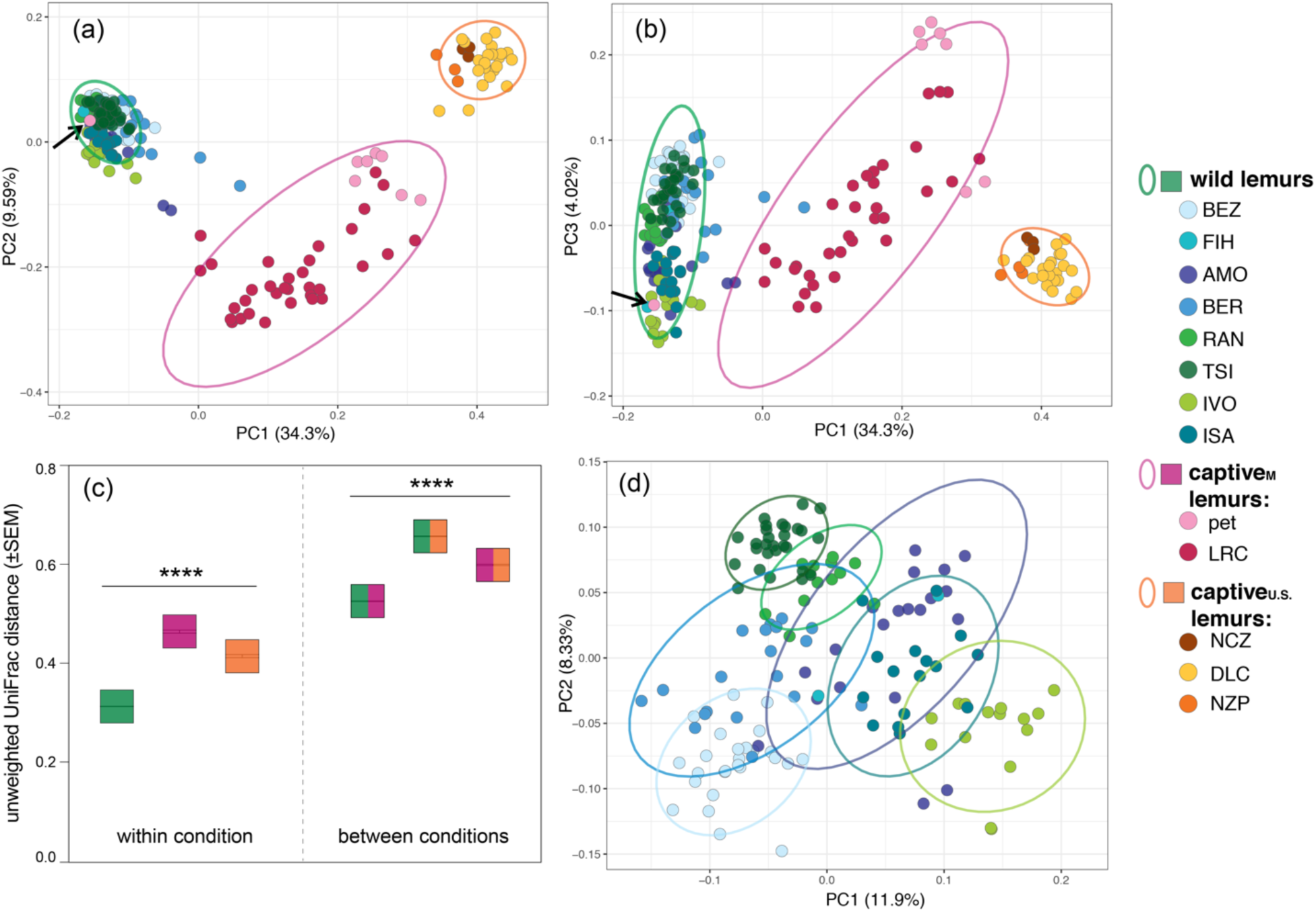
Beta diversity (unweighted UniFrac distances) of lemur gut microbiota across three environmental conditions – wilderness (wild lemurs; green), captivity in Madagascar (captive_M_ lemurs; pink), and captivity in the U.S. (captive_U.S._ lemurs; orange) – that encompass 13 setting (see Table 1 for names of abbreviated study sites). (a, b) Principal coordinate plots, showing axes 1 and 2, or 1 and 3, respectively, of individual gut microbial communities colored by setting and encircled by normal data ellipses reflecting environmental condition. (c) Mean beta diversity distance scores within (single color) and between (two colors) environmental conditions. (d) Principal coordinate plots, showing axes 1 and 2, for the eight settings within the wilderness condition. Kruskal-Wallis test with Benjamini-Hochberg correction; **** p < 0.0001.

Across the three environmental conditions, Random Forest Analysis accurately assigned 208 of the 209 gut microbial profiles to the correct condition, with a low (0.48%) out-of-bag (OOB) error rate. Based on its gut microbiota, only the previously mentioned pet lemur (see arrows in Figure 2a, b) was misclassified as a wild lemur. Across the 13 settings, Random Forest Analysis accurately classified 189 of the 209 microbial profiles (OOB error = 9.57%). The gut microbial communities of wild and captive lemurs in Madagascar were misclassified at rates of 7.9% and 7.3%, respectively, whereas those of captive lemurs in the U.S. were misclassified at a rate of 20.6%.

With respect to uniformity within environmental condition, the composition of gut microbial communities were least dissimilar between wild lemurs and most dissimilar between captive_M_ lemurs (Kruskal-Wallis test; main effect of condition on beta diversity: χ^2^ = 27487, p < 0.0001; pairwise Wilcoxon test; within wild vs. within captive_M_ lemurs: p < 0.001; within wild vs. within captive_U.S._ lemurs: p < 0.0001; Figure 4c). Between conditions, the microbiota of wild and captive_M_ lemurs were the least dissimilar, whereas the microbiota of wild vs. captive_U.S._ lemurs were the most dissimilar (pairwise Wilcoxon test: ‘wild vs. captive_M_’ vs. ‘wild vs. captive_U.S._’, p < 0.0001; ‘wild vs. captive_M_’ vs. ‘captive_M_ vs. captive_U.S._’, p < 0.0001; Figure 4c). Considering wild lemurs only, microbiota composition clustered by setting (Figure 4d). Although there was some overlap between settings, the patterns are suggestive of microbial ‘signatures’ across different settings.

### Soil microbiota: Variation in diversity, membership, and composition

#### Alpha diversity

Across the eight settings for which we sampled soil, the alpha diversity of soil microbiota did not vary significantly between conditions (Kruskal-Wallis test; Shannon diversity: χ^2^ = 3.3457, p = 0.187; Faith’s phylogenetic: χ^2^ = 3.433, p = 0.179; Figure 5) nor between settings (Kruskal-Wallis test; Shannon diversity: χ^2^ = 7.496, p = 0.379; Faith’s phylogenetic: χ^2^ = 8.936, p = 0.257; Figure 5). These null findings may owe to small sample sizes.

**Figure 5.**
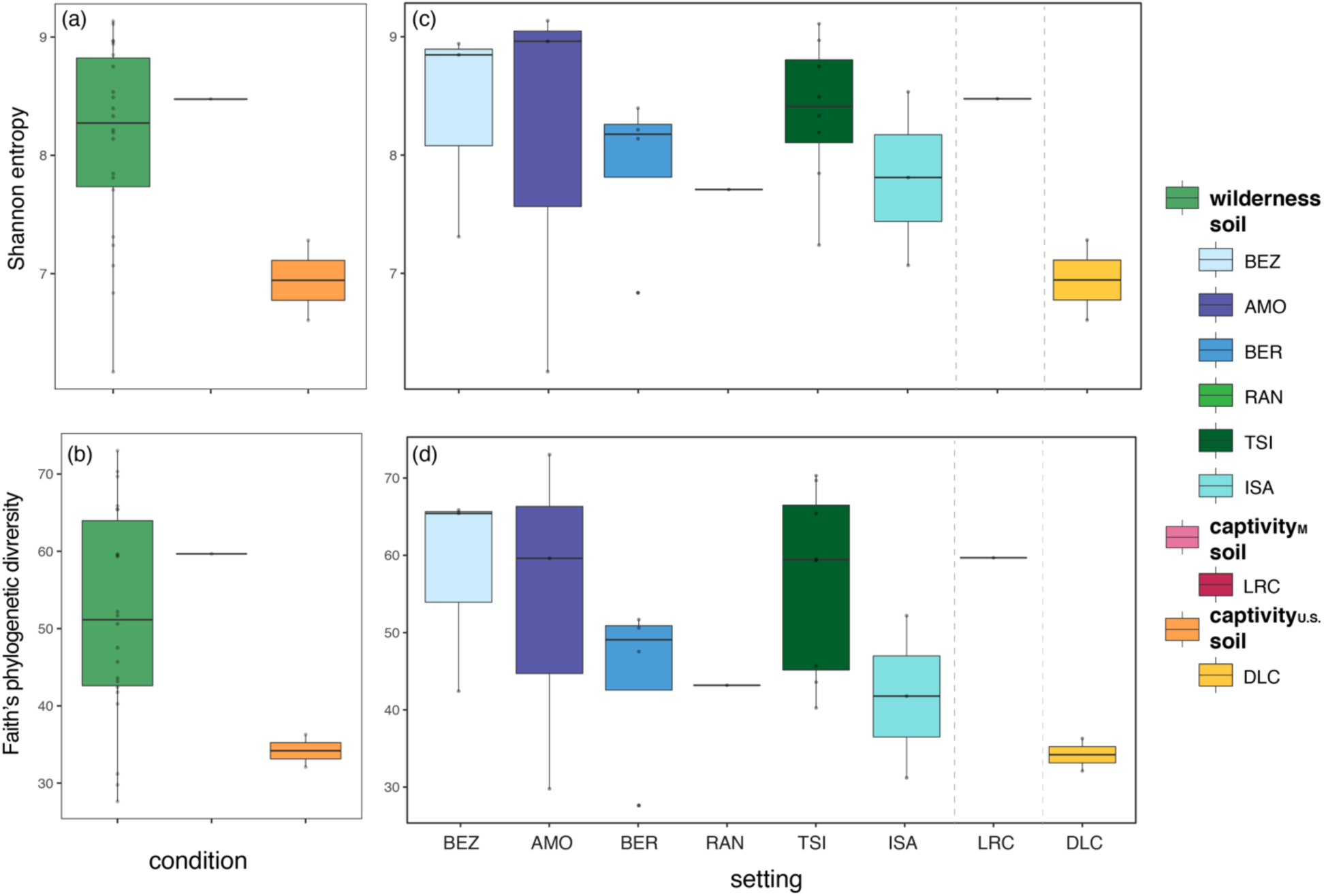
Alpha diversity metrics of soil microbiota (a, b) collapsed by environmental condition, including wilderness (wilderness soil; green), captivity in Madagascar (captivity_M_ soil; pink), and captivity in the U.S. (captivity_U.S._ soil; orange) and (c, d) averaged across individuals for each of the eight different settings (reprising the color codes of each condition, delineated by dashed vertical lines). Shown are both (a, c) Shannon diversity and (b, d) Faith’s phylogenetic diversity. Across the (c, d) settings within a condition (see Table 1 for names of abbreviated study sites), the data are plotted in descending order of mean Shannon diversity.

#### Community membership

The membership of soil communities included 77 abundant taxa, of which none were shared across all settings (Figure 6). Of the identified soil microbiota, 78.12% were unique to the soil samples and were not found in any lemur fecal samples. For the five wild populations for which we sampled soil, only five abundant taxa were shared: the genera *Bacillus* (phylum Firmicutes), *Steroidobacter* (Proteobacteria), *Bryobacter* (Acidobacteria), and *RB41* (Acidobacteria), and an unidentified member of the class Subgroup 6 (Acidobacteria). ANCOM identified nine ASVs that were differentially abundant across all soil samples, five of which (55.6%) belonged to the Balneolaceae family. In addition, compared to soil from Madagascar, the soil communities from the U.S. (hereafter, captivity_U.S._ soil) were differentially enriched for the genus *Bacillus*. By contrast, members of the family Nitrososphaeraceae (Thaumarchaeota) and the genus *Acinetobacter* (Proteobacteria) characterized soil from the wilderness and captivity settings in Madagascar (hereafter, wilderness and captivity_M_ soils, respectively; Figure S1).

**Figure 6.**
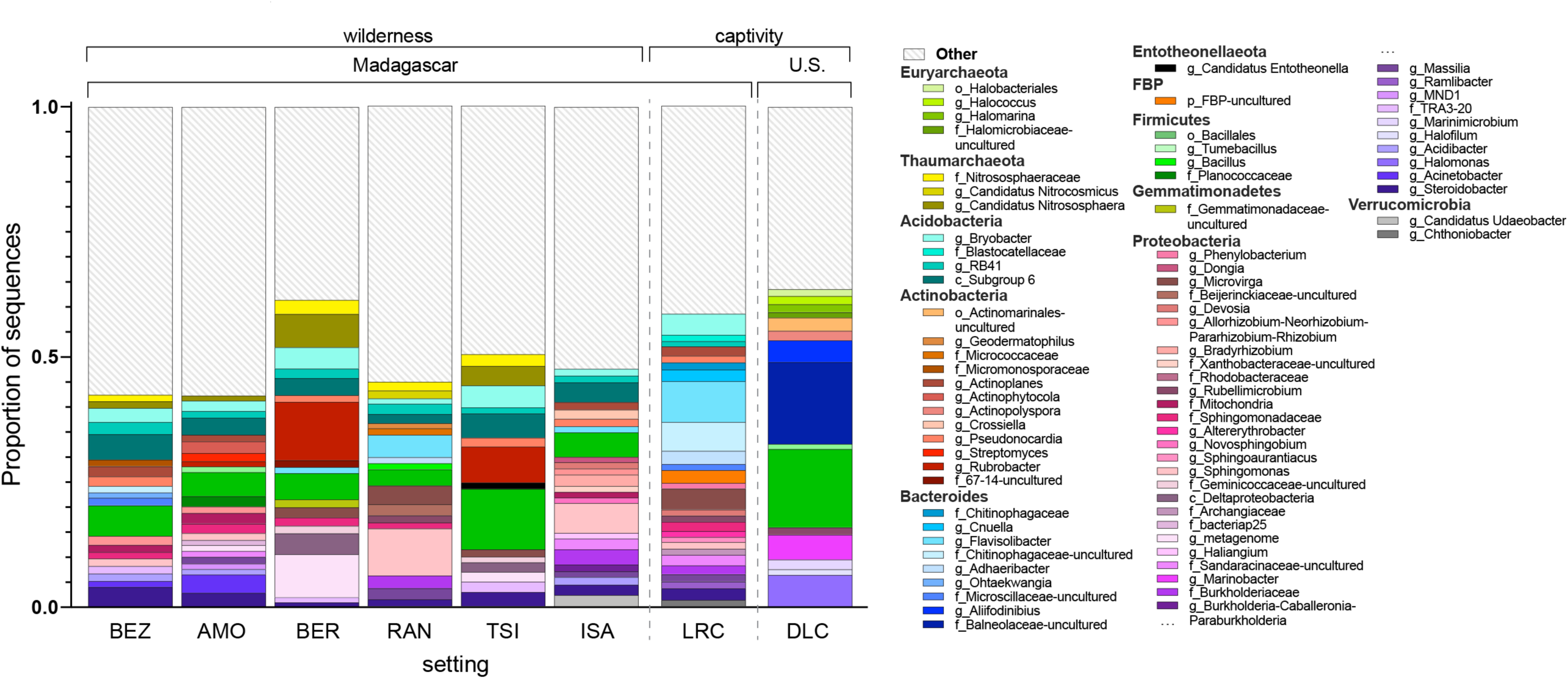
Mean proportion of sequences assigned to microbial taxa of soil at each of the eight settings sampled, within the three conditions: wilderness (wilderness soil; green), captivity in Madagascar (captivity_M_ soil; pink), and captivity in the U.S. (captivity_U.S._ soil; orange), which are delineated by dashed vertical lines (see Table 1 for names of abbreviated study sites). Taxa are identified by phylum and deepest possible taxonomic level (i.e., genus level or above); those representing < 1% of the microbiomes were combined into the category “Other”.

#### Beta diversity

The beta diversity of the soil microbiota varied between conditions (Figure 7), but only significantly so between wilderness and captivity_M_ soils (PERMANOVA; wilderness vs. captivity_M_ soils: pseudo-F = 1.337, p = 0.202; wilderness vs. captivity_U.S_ soils: pseudo-F = 3.897, p = 0.012; captivity_M_ vs. captivity_U.S_ soils: pseudo-F = 7.752, p = 0.329). Variation in soil communities within a condition was not significantly different between wilderness soils and captivity_U.S._ soils (pairwise Wilcoxon test, p = 0.130; Figure 7c). Between conditions, wilderness and captivity_M_ soils had the lowest dissimilarities (pairwise Wilcoxon test; ‘wilderness vs. captivity_M_’ vs. ‘wilderness vs. captivity_U.S_’ soils: p < 0.001; ‘wilderness vs. captivity_M_’ vs. ‘captivity_M_ vs. captivity_U.S_’: p = 0.016; ‘wild vs. captivity_U.S_’ vs. ‘captivity_M_ vs. captivity_U.S_’: p = 0.338 Figure 7c).

**Figure 7.**
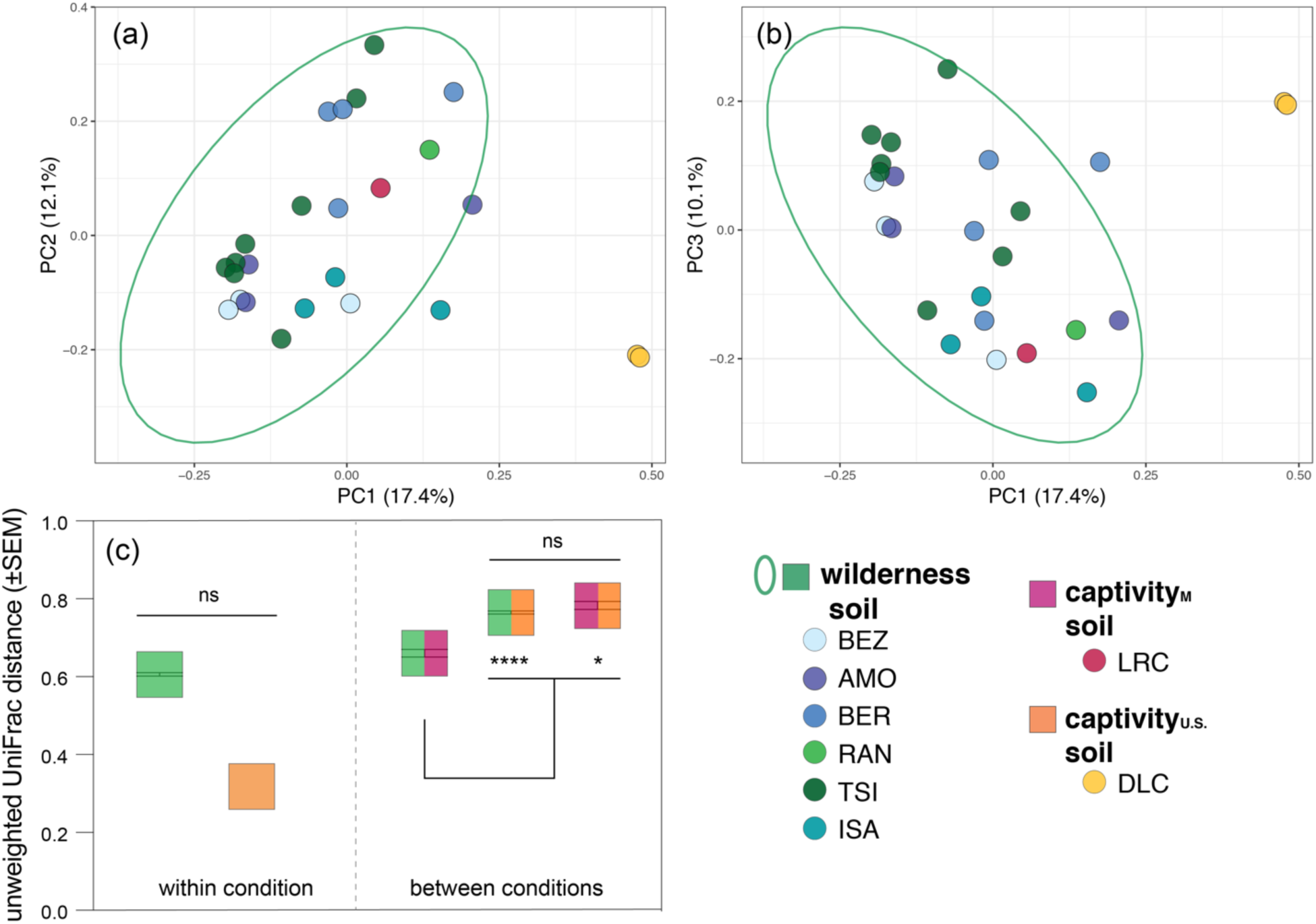
Beta diversity (unweighted UniFrac distances) of soil microbiota across three environmental conditions - wilderness (wilderness soil; green), captivity in Madagascar (captivity_M_ soil; pink), and captivity in the U.S. (captivity_U.S._ soil; orange) – that encompass eight setting (see Table 1 for names of abbreviated study sites). (a, b) Principal coordinate plots, showing axes 1 and 2, or 1 and 3, respectively, of soil microbial communities colored by setting and encircled by normal data ellipses reflecting environmental condition. (c) Mean beta diversity distance scores within (single color) and between (two colors) environmental conditions. Kruskal-Wallis test with Benjamini-Hochberg correction; * p < 0.05, **** p < 0.0001.

### Covariation of lemur gut and soil microbiota

There were 191 ASVs shared between lemur gut and soil microbiota. These were dominated by members of the Firmicutes (75 ASVs or 39.3%), Proteobacteria (49 ASVs or 25.6%), and Bacteroidetes (38 ASVs or 19.9%) phyla. Although many of the shared taxa were abundant (>1%) in either lemur gut or soil microbiota, only one genus, *Acinetobacter* (Proteobacteria), was abundant in both lemur gut and soil microbiota.

As would be predicted if environmental acquisition impacts host microbial communities, there was a significant correlation between the abundances of microbes in lemur feces and soil samples (Mantel test; r = 0.494, p < 0.001). The proportion of ‘soil-associated’ microbes found in lemur gut microbiota varied significantly across conditions (Kruskal-Wallis test; χ^2^ = 73.862, p < 0.001; Figure 8a) and settings (Kruskal-Wallis test; χ^2^ = 112.69, p < 0.001; Figure 8b). Overall, the gut microbiota of wild lemurs had significantly greater proportions of soil-associated microbes compared to those of all captive lemurs (pairwise Wilcoxon test, p < 0.001; Figure 8). In addition, captive_M_ lemurs had significantly greater proportions of soil-associated microbes in their gut microbiota compared to captive_U.S._ lemurs (pairwise Wilcoxon test; p < 0.001; Figure 8). For lemurs housed at the DLC, those that semi-free-ranged in outdoor, natural habitat enclosures had significantly greater proportions of soil-associated microbes in their gut microbiota compared to lemurs that did not have access to forested enclosures (Kruskal-Wallis test; χ^2^ = 4.641, p = 0.031; Figure 8c).

**Figure 8.**
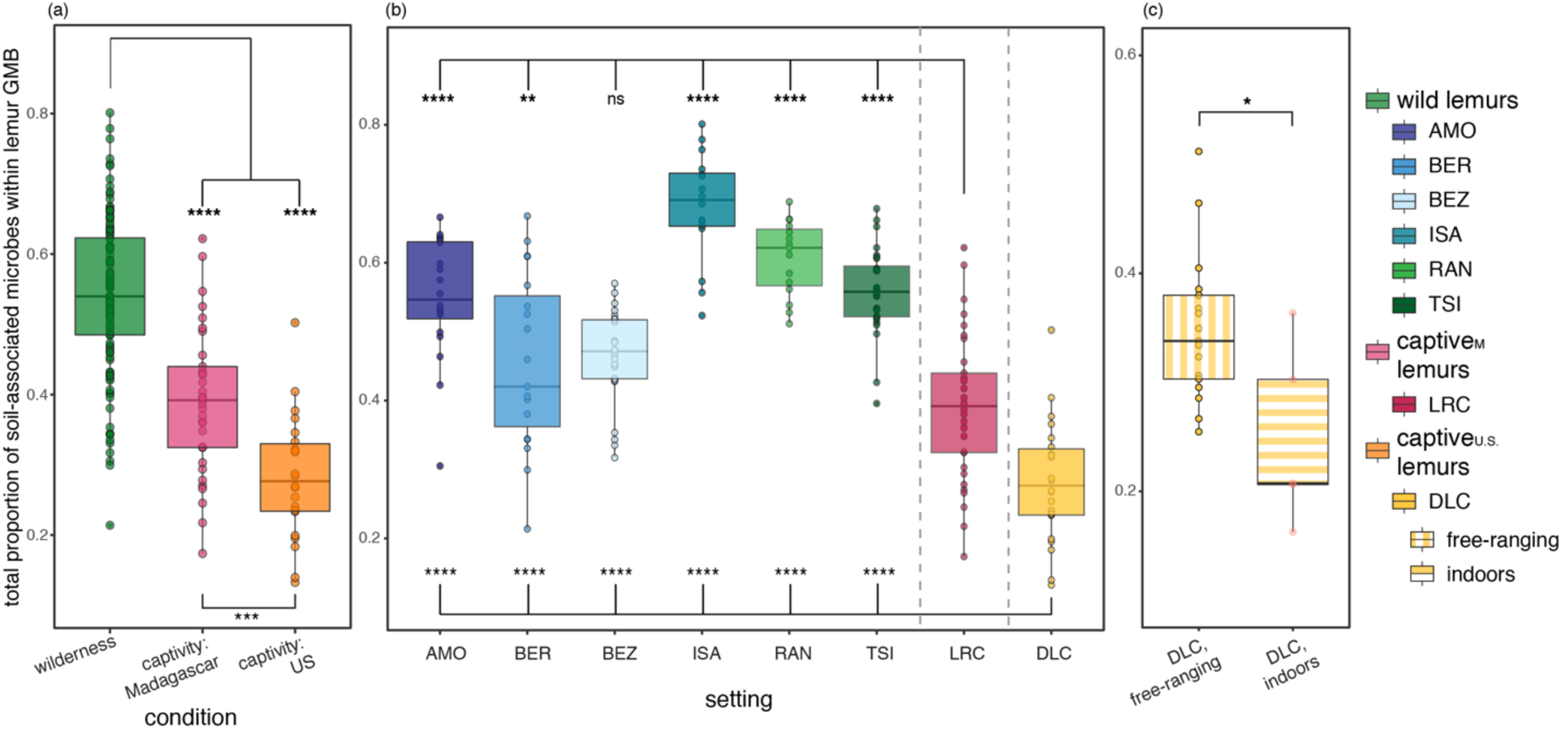
Mean proportion of total soil-associated microbes in the gut microbiota of lemurs (a) collapsed by environmental condition: wilderness (wild lemurs; green), captivity in Madagascar (captive_M_ lemurs; pink), and captivity in the U.S. (captive_U.S._ lemurs; orange). Averaged soil-associated microbes across individuals for (b) each of the eight different settings (reprising the color codes of each condition, delineated by dashed vertical lines) and (c) at the Duke Lemur Center (DLC) that were semi-free-ranging in natural habitat enclosures or were housed indoors. Kruskal-Wallis test with pairwise comparisons and Benjamini-Hochberg correction; * p < 0.05, ** p < 0.01, *** p < 0.001, **** p < 0.0001, ns = nonsignificant.

Soil from within a lemur’s setting accounted for, on average, significantly greater proportions of the lemur’s gut microbiota than did soil communities from other settings (Figure 9, Supplementary Material 2). Overall, the greatest proportion of soil-associated microbes within lemur gut microbiota occurred when the lemurs and soil were both from the wilderness (Figure 9; Supplementary Material 2). The proportion of soil-associated microbes in lemur gut microbiota dropped to near zero when comparing the lemur gut and soil microbiota between samples from the wilderness and those from captivity in the U.S. (Figure 9; Supplementary Material 2).

**Figure 9.**
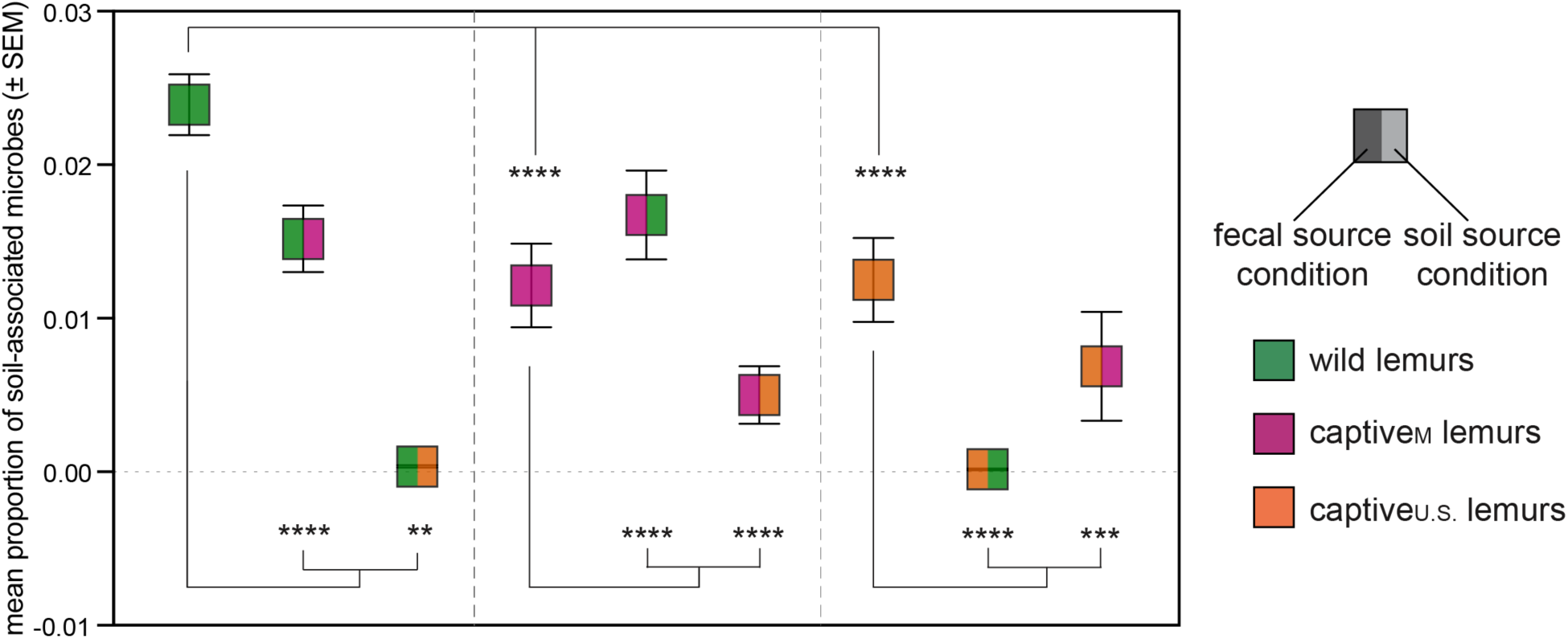
Mean proportion of soil-associated microbes in the gut microbiota of lemurs within (single color) and between (two colors) the three conditions: wilderness (wild lemurs; green), captivity in Madagascar (captive_M_ lemurs; pink), and captivity in the U.S. (captive_U.S._ lemurs; orange) – that encompass eight setting. Within the gut microbiota of lemurs from a given condition (first color = fecal source condition), values show the proportion of soil associated microbes from a given condition (second color = soil source condition). Kruskal-Wallis test with pairwise comparisons and Benjamini-Hochberg correction; ** p < 0.01, *** p < 0.001, **** p < 0.0001.

## Discussion

Through extensive fecal and soil sampling from multiple settings representing the ring-tailed lemurs’ natural range in Madagascar and in captivity on two continents, we have highlighted (1) the wide and often underrepresented variety of gut microbiota present within a single host species, (2) the lack of a universal ‘signal of captivity’ that uniformly decreases microbial diversity, (3) aspects of microbiota membership and composition that differ markedly between wild and captive populations, and (4) covariance between lemur gut and soil microbiota, which points to a key role of environmental microbes. Researchers have reported host ‘group signatures’ in microbiota, often attributed to the social transmission of microbes^5,55–58^; our results expand this concept to ‘population signatures’ and draw attention to the potential role of environmental acquisition of microbes in mediating significant inter-population variation.

Across populations of wild lemurs, we first observed substantial variation in gut microbiota diversity, membership, and composition, indicating that there is not a single ‘representative’ gut community for wild ring-tailed lemurs, as is likely the case for most host species. Nonetheless, the pattern of natural variation observed did not always meet expectations. For example, lemurs living in what is considered a relatively ‘pristine’ site, IVO – a recently discovered humid forest patch that is relatively undisturbed by human activity – unexpectedly had the second-lowest diversity of gut microbes. To the extent that lack of disturbance is a proxy of habitat quality, this pattern would be inconsistent with previous reports that greater habitat quality promotes more diverse gut microbiota^59,60^. In prior studies, the gut microbiota of ring-tailed lemurs were relatively unaffected by habitat degradation^52^. Therefore, either pristine habitats can be of low quality or the ecological and dietary flexibility of this species may dampen the impact of variation in habitat quality and type, relative to more specialized primates (e.g., folivores)^26,61–63^. That we found significant, natural, inter-population variation in a relatively hardy and robust species^49,64^ suggests that hosts with greater sensitivity to environmental variation, including habitat quality and type, would likely show even greater variation than that described herein. If so, studies constrained to single or few host populations are likely to underrepresent the wide-scale, natural variation in host gut microbiota.

We next observed gut microbiota to be compositionally distinct across populations of captive lemurs. Contrary to many previous studies^65–68^, but consistent with others^69–72^, the gut microbiota of captive lemurs were not consistently less diverse than those of wild lemurs nor were they compositionally homogenized by the similar commercial diets provided to captive subjects^67,73^. Heterogeneous gut microbiota could reflect slight differences in the diets provided (as the produce and browse available differ between captivity settings), but such minor dietary variation is unlikely to be the sole driver of such marked microbial differences, particularly in an omnivorous host. Non-dietary factors must have contributed to distinguishing the gut communities of captive lemurs. Indeed, the gut microbiota of captive lemurs in Madagascar were compositionally more similar to those of their wild counterparts than to those of captive lemurs in the U.S. Based on this observation, we suggest that the effect of a commercial diet is not necessarily the strongest differentiator of gut consortia and that the effects of captivity cannot be standardized across populations. Our results raise questions about the commonly held view that greater alpha diversity is both (a) a hallmark of wild individuals and (b) a proxy for a healthier gut community^74–78^. Although we did not assess gut health, pet lemurs fed diets of rice and fruit, living in close contact with people and domestic animals, often housed solitarily indoors, are prone to disease^79–81^; yet, their gut consortia were as diverse as those of wild lemurs living at the relatively pristine site, IVO. Moreover, captive lemurs from the DLC and NCZ in the U.S. had some of the most diverse gut consortia, equaling the greatest diversity seen in wild lemurs (e.g., in BEZ lemurs). These results add to the mounting evidence^61,82,83^ that alpha diversity alone, without the context of host ecology and other microbiome data, should not be used to extrapolate about the health state of gut consortia or the host’s environment.

We also found that, between wild and captive lemurs, the membership and composition of gut microbiota was indicative of the environmental condition. There was little evidence of a diverse ‘core’ microbiome, as only four taxa were found to be abundant across all lemur populations. Two of those core taxa, *Erysipelotrichaceae UCG-004* and *Treponema 2*, were differentially abundant between the three conditions. Despite links between members of *Erysipelotrichaceae* and high-fat, commercial diets in humans^84^, *Erysipelotrichaceae* microbes were reported to be enriched in wild compared to captive chimpanzees^85^, mirroring our findings in lemurs. Furthermore, the genus *Erysipelotrichaceae UCG-004* was more abundant in the gut microbiota of chimpanzees, relative to humans^86^, and in folivorous woolly lemurs compared to other lemur species^87^. The functionally diverse members of the *Treponema* genus were more abundant in the gut microbiota of captive vs. wild hosts in other species^85,88^. *Treponema* members break down pectin^89,90^, a complex plant polysaccharide enriched in ripe fruits, such as those commonly provided to captive ring-tailed lemurs^91,92^. Compositionally, the gut microbiota of wild lemurs were markedly less varied than those of lemurs in all captivity settings. These findings support the “Anna Karenina” principle^93,94^, which posits that perturbations of microbiota result in unstable communities and, thus, ‘unperturbed’ hosts have less variation in their microbiota than do ‘disturbed’ hosts. A single exception to the gut microbiota clustering according to the hosts’ conditions was a pet lemur with gut microbiota that resembled that of wild lemurs. Although we can only speculate about this individual’s history, if recently taken from their natural habitat, the gut microbiota could still reflect the wild origins of this animal, potentially indicative of gradual change in response to environmental shifts^95,96^.

Lastly, we observed that patterns in lemur gut microbiota were somewhat mirrored in the diversity and composition of soil microbiota, suggesting that environmental conditions other than diet, including exposure to external microbes in soils, may influence gut microbiomes^97^. Madagascar’s geographical isolation for ∼88 million years accounts for high levels of floral and faunal endemism^98,99^. The same is true of microbes, as evidenced by the numerous, unique pathogenic microorganisms found on the island^100–103^. Unsurprisingly, therefore, soil microbiota in Madagascar, whether originating in wilderness or captivity settings, were similar in composition and significantly divergent from soils in the U.S.^104^. Given the disparate geographic distributions of many wild vs. captive animals, environmental acquisition that reflects local microbial endemism may be particularly relevant for distinguishing gut microbiota between wild and captive conspecifics. For example, the natural ranges of most primates occur in the tropics^105,106^, yet most accredited zoos and captive facilities that house primates are found outside of tropical regions (in e.g., Europe and North America)^107,108^; the distinct environmental consortia surrounding wild and captive conspecifics should reflect their geographic or continental divides.

Regarding the exposure to environmental microbes, soil-associated microbes were more prevalent in lemurs that had greater exposure to natural environments and the acquired soil microbes were specific to the lemurs’ environment, reflecting active environmental acquisition. This observation expands on findings that abiotic soil properties mediate primate gut microbiota^97^. Wild and captive ring-tailed lemurs perform geophagy (i.e., earth-eating), a behavior that is linked to nutrient and microbial supplementation^109,110^ and is a potential vector for the incorporation of environmental microbes^39^. Similarly, dietary items may act as vessels of soil or environmental microbes^40^; dietary variation across wild and captive lemurs may influence gut microbiomes by simultaneously offering different nutrients and different microbes. Akin to most cross-sectional studies of microbiomes, we were unable to assess the persistence or viability of the soil-associated microbes in lemur gut communities. It is, therefore, possible that the soil-associated microbes in lemur guts were ephemeral or non-viable; however, our results indicate setting-specific, environmental acquisition, supporting that these patterns are not random and that the acquired microbes may be subject to filters that enable the incorporation of only specific microbes^20,111,112^. Furthermore, we analyzed these data from the perspective that environmental consortia act as sources of microbes for host-associated communities, but we expect consistent, bidirectional transmission of microbes between hosts and their environments, a relationship that warrants further investigation.

While expanding our understanding of the factors that shape host-microbe relationships, these results also have significant potential to inform animal care and conservation strategies. Perturbed microbiota are increasingly recognized as culprits of obesity, gastrointestinal distress, and even associated mortality in captive animals^75,113–115^. Given that lemurs are among the most endangered mammals on the planet^116^, maintaining populations of healthy animals in captivity is an important ‘safety net’ that augments *in-vivo* conservation efforts^117,118^. We suggest that environmental acquisition may be a key component of ‘rewilding’ or ‘bioaugmenting’ captive animal gut microbiota, a process by which gut consortia can be reshaped to better promote host-microbe symbiosis^26,117,119^. Identifying what comprises healthy gut microbiota is a complex and ongoing area of research; nonetheless, we show that environmental acquisition is a potential driver of microbial communities and thus should be considered as a component of animal health.

## Conclusions

Even in a relatively robust, omnivorous host, gut microbiota are distinct across populations. This variation reflects environmental variability that is underrepresented by a simple wild vs. captive dichotomy. Moreover, concurrent analysis of lemur gut and soil microbiota supports the premise that environmental acquisition contributes to shaping host-associated microbiota; hosts and their associated microbes are components of a larger landscape that includes interactions with environmental microbes. Together, these results expand our understanding of intraspecific host-microbe dynamics under varying environmental conditions and reinforce the value of broad-scale, comparative investigations of microbial variation within a single host species.

## Methods

### Study sites

Our research sites included 13 settings (one per ‘population’), grouped under the following three environmental conditions: wilderness in Madagascar (8 settings), captivity in Madagascar (2 settings), and captivity in the U.S. (3 settings; Table 1). The wilderness settings occurred in protected areas (e.g., national parks, community-managed reserves) that varied in habitat type (Table 1). The captivity settings in Madagascar included the Lemur Rescue Center (LRC; Toliara, Madagascar), where the animals were socially housed, and various townships that were home to individual pets. Lastly, the captivity settings in the U.S. included the North Carolina Zoo (NCZ; Asheboro, NC), the Duke Lemur Center (DLC; Durham, NC), and the National Zoological Park (NZP; Washington, DC). These facilities were comparable to one another, all with socially housed lemurs.

### Subjects

Across all research sites, our subjects included 215 adult, ring-tailed lemurs (82 male, 81 female, 52 of unknown sex; Table 1). The wilderness sites were each occupied by multiple lemur troops, ranging in size from 5-24 individuals. Excluding the pets, all captive settings included groups of 2-7 lemurs that had access to indoor and outdoor enclosures, and were provided facility-standardized diets (i.e., fresh produce and commercial chow, freely available water). Certain animals at the LRC and the DLC also had access to natural habitat enclosures that, respectively, consisted of dry and spiny forest (LRC) or North American deciduous and pine hardwood forest (DLC). The pets were kept in human dwellings (i.e., houses or hotels) and were fed fruit, rice, and other foods intended for human consumption.

### Sample collection

During a span of four years (2016-2020), we collected ‘matched’ fecal and soil samples from our subjects and study sites, respectively. Within 8 weeks of fecal or soil collection, the samples were transported to the U.S., where they were stored at −80 °C, until analysis.

For feces, we opportunistically collected fresh samples, upon the lemur’s observed voiding. In Madagascar, collections occurred during the dry season (May-October) and, in the U.S., collections occurred end of summer through fall (August-November). To avoid soil contamination of the fecal sample, we removed the outer layer of each fecal pellet. We then placed the sample in an Omnigene tube that contained a stabilizing buffer that preserved microbial communities at room temperature for 8 weeks (Omnigene.Gut tube, DNAgenotek, Ontario, Canada^120,121^). All settings were represented by fecal samples from minimally two lemurs (the maximum number of individuals represented was 33).

When collecting soil in nature, we avoided high-defecation areas (e.g., under sleeping trees) while identifying core areas where lemurs most commonly spent time on the ground. Within these core areas, we demarcated a 2-3 m^2^ area and collected soil from each of five evenly spaced locations, using a clean, individually wrapped, sterile plastic spatula. For each area, the five aliquots of topsoil (top 2-3 cm of soil) were pooled in a single Omnigene tube to create a representative soil sample. Because multiple lemur troops inhabited each of the wilderness settings, in some cases with overlapping core areas, we prioritized collecting soil samples from areas of maximal use. In some cases, we were unable to collect soil samples for every troop that provided fecal samples. At the LRC and DLC, we used the same collection methods to collect soil samples from areas in the natural habitat enclosures where lemurs semi-free-ranged. Because it is illegal to own pet lemurs in Madagascar, we minimized owner concern by collecting only fecal samples for this group. Because of other logistical and analytical constraints (see below), only eight of the 13 settings were represented by usable, pooled soil samples.

### Microbial DNA extraction and sequencing

Following the manufacturer’s protocols for the DNeasy Powersoil kit (QIAGAN, Frederick, MD), we extracted bacterial genomic DNA from fecal and soil samples. We quantified DNA using a Fluorometer (broad-spectrum kit, Qubit 4, Thermo Fisher Scientific, Waltham, MA). Aliquots of extracted DNA were sent to Argonne National Laboratory’s Environmental Sequencing facility (Lemont, IL) for library preparation and amplicon sequencing of the 16S rRNA gene. After amplification of the V4 region with region-specific primers and sample-specific 12-base barcodes, samples were pooled and amplicon libraries were cleaned using AMPure XP Beads. Amplicons were then sequenced on a 151 x 151 base pair Illumina MiSeq run^122^.

### Bioinformatics and statistics

We processed the raw sequence data using a previously published bioinformatics pipeline generated in QIIME2^123^. In brief, we used the pipeline to join forward and reverse reads, demultiplex and quality filter the joined reads (DADA2), generate a phylogenetic tree, and assign taxonomy based on 99% sequence similarity (SILVA database ^124,125^, ver. 138.1). After quality filtering, samples with fewer than 10,000 sequences were removed from downstream analyses, resulting in 209 fecal samples and 25 soil samples with over 11 million combined reads and an average of ∼50,000 reads per sample. To visually represent rare taxa that had relative abundances < 1% of the total sequences, we combined them into the conglomerate “Other” category (Figures 1 and 6). Using tables of amplicon sequence variants (ASVs), we calculated metrics of alpha diversity (Shannon and Faith’s Phylogenetic diversity metric) and beta diversity (UniFrac distances).

To test for differences in alpha diversity between the gut microbiota of lemurs under the three environmental conditions and in the 13 settings, we first used generalized linear models (GLMs; glm in R, ver, 4.0.2) with condition or setting and sex as fixed effects. To further test for variation in lemur gut microbiota and soil microbiota alpha diversity, we used nonparametric statistics (e.g., Kruskal-Wallis tests, and pairwise Wilcoxon rank sum tests with Benjamini-Hochberg adjustment) to perform pairwise comparisons between the various conditions and settings. To identify and test for effects of condition or setting on beta diversity (unweighted UniFrac distances) in lemur fecal and soil microbiota, we used principal coordinate analysis (i.e., to visualize clustering of microbiota composition) and Permutational Multivariate Analysis of Variance (PERMANOVA) in QIIME2. We then performed Random Forest Analysis^126^, which is a supervised learning technique that uses decision trees to classify data to specific categories and provides an overall model error rate (out of the bag error or OOB error). To identify microbes enriched in specific groups of samples, we used differential abundance analyses via Analysis of Compositions of Microbiomes (ANCOM) and songbird software^127^ in QIIME2, paired with visualization through Qurro^128^.

For the eight settings where we obtained matched fecal and soil samples (Table 1), we analyzed covariation between lemur gut microbiota and the associated soil communities by performing a Mantel test on microbial abundance matrices of lemur gut and soil microbiota. Because multiple lemur fecal samples were associated with each soil sample, we created comparable matrices for the Mantel test by averaging the microbial abundances across the fecal samples of lemurs directly associated with a given soil sample, resulting in a single, mean lemur gut community associated with each soil community. For this process, we omitted fecal samples from troops not represented by a soil sample or for which troop identity was unknown.

To test if soil-associated microbes were present in lemur gut microbiota, we used FEAST, a tool for fast expectation-maximization microbial source tracking^129^. FEAST assumes each ‘sink’ sample is a convex combination of known and unknown ‘sources’ and uses multinomial distributions and machine-learning classification to model the microbial source-sink data^129^. For this analysis, we used the matched lemur gut and soil samples; all soil samples collected in a given setting were used to represent the potential exposure to environmental microbes experienced by all sampled lemurs in that same setting, regardless of troop identity. Because we were testing whether environmental acquisition influences lemur gut microbiota, and because this analysis requires an assumption of directionality (i.e., from a source to a sink), we categorized soil samples as ‘sources’ and lemur fecal samples as ‘sinks’; however, we acknowledge and discuss the potential for bi-directional transmission of microbes between lemurs and soil. For each lemur fecal sample, we calculated the proportions of microbes that were identified as stemming from each soil community and from a default ‘unknown source’ that accounts for microbes not relevant to soil microbiota. Lastly, we used FEAST to test for differences in the proportion of soil microbes in the gut microbiota of lemurs at the DLC that were either semi-free-ranging or sequestered to indoor enclosures.

## Declarations

### Ethics approval and consent to participate

Sampling in Madagascar occurred with approval from Madagascar National Parks and appropriate governmental agencies (Ministry of Environment, Ecology, and Forests; permit #s 147/18/MEEF/SG/DGF/DSAP/SCB.Re, 152/19/MEDD/SG/DGEF/DGRNE, 159/16/MEEF/SG/DGF/DSAP/SCB.Re, 154/17/ MEEF/SG/DGF/DSAP/SCB.Re, 156/19/MEEF/SG/DGF/DSAP/SCB.Re). Sampling at the DLC, NCZ, and NZP occurred with approval from the appropriate Animal Care and Use Committees (Duke University’s Institutional Animal Care and Use Committee: protocol #A111-16-05; North Carolina Zoo Animal Care and Use Committee: approved without protocol number; Smithsonian National Zoological Park Research Animal Care and Use Committee: approved without protocol number).

### Consent for publication

Not applicable. This study does not contain any individual person’s data in any form.

### Availability of data and material

Sequencing reads are available in the National Center for Biotechnology Information’s Sequence Read Archive (BioProject ID #TBD, BioSample accession #s TBD). Additional datasets generated and/or analyzed during the current study are available from the corresponding author upon reasonable request.

### Competing interests

We attest that no author has financial or non-financial competing interests.

### Funding

Funding was provided by awards from the National Science Foundation (BCS 1749465 to CMD), the Triangle Center for Evolutionary Medicine (Graduate Student Research Award to SLB), the Kenan Institute for Ethics at Duke University (Anthropocene Graduate Research Grant to SLB). During collections, ML was funded by the Margot Marsh Biodiversity Fund.

### Authors’ contributions

SLB and CMD conceived of the study, with input from LKG. SLB, LKG, SR, SC, RSR, TAC, and ML collected samples, documented metadata, and transported materials/samples. SLB performed the bioinformatic and statistical analyses. SLB and CMD wrote the manuscript and all authors read and approved the submitted version.

## Acknowledgements

For their assistance with sample collection in the wilderness sites, we are deeply grateful to Laurent ‘Raleso’ Randrianasolo, Remi Rakotovao, Georges René Rakotonirina, Soatata Honore Reseva, Chelsea Southworth, Melina Nolas, and Lauren Petronaci. We thank Dr. Patricia Wright and the Centre ValBio for providing accessibility to the IVO collection site and assistance in sample transportation and storage. We further thank the current and past staff members of the LRC, DLC, NCZ, and NZP for their assistance with sample collection in captivity settings. We are grateful to Sarah Owens at Argonne National Laboratory for providing guidance and sequencing services. This is DLC publication number (##TBD).

## Notes

### Competing Interest Statement

The authors have declared no competing interest.

### Summary of Updates

This version of the manuscript has been revised to include the correct the author line in the downloadable PDF.

## References

1. Hansen J, Gulati A, Sartor RB. The role of mucosal immunity and host genetics in defining intestinal commensal bacteria. Curr Opin Gastroenterol 2010; 26: 564.

2. Amato KR, Sanders JG, Song SJ, Nute M, Metcalf JL, Thompson LR, Morton JT, Amir A, McKenzie VJ, Humphrey G. Evolutionary trends in host physiology outweigh dietary niche in structuring primate gut microbiomes. ISME J 2019; : 1.

3. Milani C, Alessandri G, Mancabelli L, Mangifesta M, Lugli GA, Viappiani A, Longhi G, Anzalone R, Duranti S, Turroni F. Deciphering the impact of diet and host physiology on the mammalian gut microbiome by multi-omics approaches. Appl Environ Microbiol 2020.

4. Rothschild D, Weissbrod O, Barkan E, Kurilshikov A, Korem T, Zeevi D, Costea PI, Godneva A, Kalka IN, Bar N. Environment dominates over host genetics in shaping human gut microbiota. Nature 2018; 555: 210–215.

5. Tung J, Barreiro LB, Burns MB, Grenier JC, Lynch J, Grieneisen LE, Altmann J, Alberts SC, Blekhman R, Archie EA. Social networks predict gut microbiome composition in wild baboons. Elife 2015; 2015. doi:10.7554/eLife.05224.

6. Tasnim N, Abulizi N, Pither J, Hart MM, Gibson DL. Linking the gut microbial ecosystem with the environment: does gut health depend on where we live? Front Microbiol 2017; 8: 1935.

7. David LA, Maurice CF, Carmody RN, Gootenberg DB, Button JE, Wolfe BE, Ling A V, Devlin AS, Varma Y, Fischbach MA. Diet rapidly and reproducibly alters the human gut microbiome. Nature 2014; 505: 559.

8. Kartzinel TR, Hsing JC, Musili PM, Brown BRP, Pringle RM. Covariation of diet and gut microbiome in African megafauna. Proc Natl Acad Sci 2019; 116: 23588–23593.

9. Youngblut ND, Reischer GH, Walters W, Schuster N, Walzer C, Stalder G, Ley RE, Farnleitner AH. Host diet and evolutionary history explain different aspects of gut microbiome diversity among vertebrate clades. Nat Commun 2019; 10: 1–15.

10. Greene LK, McKenney EA, O’Connell TM, Drea CM. The critical role of dietary foliage in maintaining the gut microbiome and metabolome of folivorous sifakas. Sci Rep 2018; 8: 14482.

11. Peccia J, Kwan SE. Buildings, beneficial microbes, and health. Trends Microbiol 2016; 24: 595–597.

12. Hyde ER, Navas-Molina JA, Song SJ, Kueneman JG, Ackermann G, Cardona C, Humphrey G, Boyer D, Weaver T, Mendelson JR. The oral and skin microbiomes of captive komodo dragons are significantly shared with their habitat. MSystems 2016; 1: e00046–16.

13. Cardona C, Lax S, Larsen P, Stephens B, Hampton-Marcell J, Edwardson CF, Henry C, Van Bonn B, Gilbert JA. Environmental sources of bacteria differentially influence host-associated microbial dynamics. MSystems 2018; 3.

14. Kuthyar S, Manus MB, Amato KR. Leveraging non-human primates for exploring the social transmission of microbes. Curr Opin Microbiol 2019; 50: 8–14.

15. Durrer S, Schmid-Hempel P. Shared use of flowers leads to horizontal pathogen transmission. Proc R Soc London Ser B Biol Sci 1994; 258: 299–302.

16. Kulkarni S, Heeb P. Social and sexual behaviours aid transmission of bacteria in birds. Behav Processes 2007; 74: 88–92.

17. Pedersen AB, Davies TJ. Cross-species pathogen transmission and disease emergence in primates. Ecohealth 2009; 6: 496–508.

18. Arora N, Sadovsky Y, Dermody TS, Coyne CB. Microbial vertical transmission during human pregnancy. Cell Host Microbe 2017; 21: 561–567.

19. Funkhouser LJ, Bordenstein SR. Mom knows best: the universality of maternal microbial transmission. PLoS Biol 2013; 11: e1001631.

20. Walke JB, Becker MH, Loftus SC, House LL, Cormier G, Jensen R V, Belden LK. Amphibian skin may select for rare environmental microbes. ISME J 2014; 8: 2207.

21. Seedorf H, Griffin NW, Ridaura VK, Reyes A, Cheng J, Rey FE, Smith MI, Simon GM, Scheffrahn RH, Woebken D. Bacteria from diverse habitats colonize and compete in the mouse gut. Cell 2014; 159: 253–266.

22. Smith CCR, Snowberg LK, Caporaso JG, Knight R, Bolnick DI. Dietary input of microbes and host genetic variation shape among-population differences in stickleback gut microbiota. ISME J 2015; 9: 2515–2526.

23. Selway CA, Mills JG, Weinstein P, Skelly C, Yadav S, Lowe A, Breed MF, Weyrich LS. Transfer of environmental microbes to the skin and respiratory tract of humans after urban green space exposure. Environ Int 2020; 145: 106084.

24. Schmidt E, Mykytczuk N, Schulte-Hostedde AI. Effects of the captive and wild environment on diversity of the gut microbiome of deer mice (Peromyscus maniculatus). ISME J 2019; 13: 1293–1305.

25. McKenzie VJ, Song SJ, Delsuc F, Prest TL, Oliverio AM, Korpita TM, Alexiev A, Amato KR, Metcalf JL, Kowalewski M. The Effects of Captivity on the Mammalian Gut Microbiome. Integr Comp Biol 2017; 57: 690–704.

26. Greene LK, Bornbusch SL, McKenney EA, Harris RL, Gorvetzian SR, Yoder AD, Drea CM. The importance of scale in comparative microbiome research: New insights from the gut and glands of captive and wild lemurs. Am J Primatol 2019.

27. Clayton JB, Vangay P, Huang H, Ward T, Hillmann BM, Al-Ghalith GA, Travis DA, Long HT, Tuan B Van, Minh V Van et al. Captivity humanizes the primate microbiome. Proc Natl Acad Sci 2016; 113: 10376–10381.

28. Yao R, Xu L, Hu T, Chen H, Qi D, Gu X, Yang X, Yang Z, Zhu L. The “wildness” of the giant panda gut microbiome and its relevance to effective translocation. Glob Ecol Conserv 2019; 18: e00644.

29. Frankel JS, Mallott EK, Hopper LM, Ross SR, Amato KR. The effect of captivity on the primate gut microbiome varies with host dietary niche. Am J Primatol 2019; 81: e23061.

30. Bornbusch SL, Greene LK, McKenney EA, Volkoff SJ, Midani FS, Joseph G, Gerhard WA, Iloghalu U, Granek J, Gunsch CK. A comparative study of gut microbiomes in captive nocturnal strepsirrhines. Am J Primatol 2019; 81: e22986.

31. Gibson KM, Nguyen BN, Neumann LM, Miller M, Buss P, Daniels S, Ahn MJ, Crandall KA, Pukazhenthi B. Gut microbiome differences between wild and captive black rhinoceros–implications for rhino health. Sci Rep 2019; 9: 1–11.

32. Ren T, Boutin S, Humphries MM, Dantzer B, Gorrell JC, Coltman DW, McAdam AG, Wu M. Seasonal, spatial, and maternal effects on gut microbiome in wild red squirrels. Microbiome 2017; 5: 1–14.

33. Hicks AL, Lee KJ, Couto-Rodriguez M, Patel J, Sinha R, Guo C, Olson SH, Seimon A, Seimon TA, Ondzie AU. Gut microbiomes of wild great apes fluctuate seasonally in response to diet. Nat Commun 2018; 9: 1–18.

34. Baniel A, Amato KR, Beehner JC, Bergman TJ, Mercer A, Perlman RF, Petrullo L, Reitsema L, Sams S, Lu A. Seasonal shifts in the gut microbiome indicate plastic responses to diet in wild geladas. Microbiome 2021; 9: 1–20.

35. Guarner F, Bourdet-Sicard R, Brandtzaeg P, Gill HS, McGuirk P, Van Eden W, Versalovic J, Weinstock J V, Rook GAW. Mechanisms of disease: the hygiene hypothesis revisited. Nat Clin Pract Gastroenterol Hepatol 2006; 3: 275–284.

36. Chen C-Y, Chen P-C, Weng FC-H, Shaw GT-W, Wang D. Habitat and indigenous gut microbes contribute to the plasticity of gut microbiome in oriental river prawn during rapid environmental change. PLoS One 2017; 12: e0181427.

37. Mushegian AA, Arbore R, Walser J-C, Ebert D. Environmental sources of bacteria and genetic variation in behavior influence host-associated microbiota. Appl Environ Microbiol 2019; 85.

38. Sylvain F-É, Derome N. Vertically and horizontally transmitted microbial symbionts shape the gut microbiota ontogenesis of a skin-mucus feeding discus fish progeny. Sci Rep 2017; 7: 1–14.

39. Borruso L, Checcucci A, Torti V, Correa F, Sandri C, Luise D, Cavani L, Modesto M, Spiezio C, Mimmo T. I Like the Way You Eat It: Lemur (Indri indri) Gut Mycobiome and Geophagy. Microb Ecol 2021; : 1–9.

40. Li H, Li T, Yao M, Li J, Zhang S, Wirth S, Cao W, Lin Q, Li X. Pika gut may select for rare but diverse environmental bacteria. Front Microbiol 2016; 7: 1269.

41. Kikuchi Y, Hosokawa T, Fukatsu T. Insect-microbe mutualism without vertical transmission: a stinkbug acquires a beneficial gut symbiont from the environment every generation. Appl Environ Microbiol 2007; 73: 4308–4316.

42. Inoue R, Ushida K. Vertical and horizontal transmission of intestinal commensal bacteria in the rat model. FEMS Microbiol Ecol 2003; 46: 213–219.

43. Leftwich PT, Edgington MP, Chapman T. Transmission efficiency drives host–microbe associations. Proc R Soc B 2020; 287: 20200820.

44. Maamar S Ben, Hu J, Hartmann EM. Implications of indoor microbial ecology and evolution on antibiotic resistance. J Expo Sci Environ Epidemiol 2020; 30: 1–15.

45. Hartmann EM, Hickey R, Hsu T, Betancourt Román CM, Chen J, Schwager R, Kline J, Brown GZ, Halden RU, Huttenhower C. Antimicrobial chemicals are associated with elevated antibiotic resistance genes in the indoor dust microbiome. Environ Sci Technol 2016; 50: 9807–9815.

46. Thompson LR, Sanders JG, McDonald D, Amir A, Ladau J, Locey KJ, Prill RJ, Tripathi A, Gibbons SM, Ackermann G. A communal catalogue reveals Earth’s multiscale microbial diversity. Nature 2017; 551: 457–463.

47. Jolly A, Sussman RW, Koyama N, Rasamimanana H. Ringtailed Lemur Biology: Lemur catta in Madagascar (Google eBook). 2006 http://books.google.com/books?id=WQ5yWlEJVVYC&pgis=1.

48. Gould L. Lemur catta ecology: what we know and what we need to know. In: Lemurs. Springer, 2006, pp 255–274.

49. Gabriel DN. Habitat Use and Activity Patterns as an Indication of Fragment Quality in a Strepsirrhine Primate. Int J Primatol 2013; 34: 388–406.

50. Mason GJ. Species differences in responses to captivity: stress, welfare and the comparative method. Trends Ecol Evol 2010; 25: 713–721.

51. Bornbusch SL, Harris RL, Grebe NM, Dimac-Stohl K, Drea CM. Longitudinal effects of antibiotics and fecal transplant on lemur gut microbiota structure, associations, and resistomes. doi:https://doi.org/10.1101/2020.11.11.378349;

52. Bennett G, Malone M, Sauther ML, Cuozzo FP, White B, Nelson KE, Stumpf RM, Knight R, Leigh SR, Amato KR. Host age, social group, and habitat type influence the gut microbiota of wild ring-tailed lemurs (Lemur catta). Am J Primatol 2016; 78: 883–892.

53. Fogel AT. The gut microbiome of wild lemurs: a comparison of sympatric Lemur catta and Propithecus verreauxi. Folia Primatol 2015; 86: 85–95.

54. Dias PC. Sources and sinks in population biology. Trends Ecol Evol 1996; 11: 326–330.

55. Leclaire S, Nielsen JF, Drea CM. Bacterial communities in meerkat anal scent secretions vary with host sex, age, and group membership. Behav Ecol 2014; 25: 996–1004.

56. Theis KR, Schmidt TM, Holekamp KE. Evidence for a bacterial mechanism for group-specific social odors among hyenas. Sci Rep 2012; 2: 615.

57. Vernier CL, Chin IM, Adu-Oppong B, Krupp JJ, Levine J, Dantas G, Ben-Shahar Y. The gut microbiome defines social group membership in honey bee colonies. Sci Adv 2020; 6: eabd3431.

58. Sarkar A, Harty S, Johnson KV-A, Moeller AH, Archie EA, Schell LD, Carmody RN, Clutton-Brock TH, Dunbar RIM, Burnet PWJ. Microbial transmission in animal social networks and the social microbiome. Nat Ecol Evol 2020; 4: 1020–1035.

59. Amato KR, Yeoman CJ, Kent A, Righini N, Carbonero F, Estrada A, Gaskins HR, Stumpf RM, Yildirim S, Torralba M. Habitat degradation impacts black howler monkey (Alouatta pigra) gastrointestinal microbiomes. ISME J 2013; 7: 1344.

60. Barelli C, Albanese D, Donati C, Pindo M, Dallago C, Rovero F, Cavalieri D, Tuohy KM, Hauffe HC, De Filippo C. Habitat fragmentation is associated to gut microbiota diversity of an endangered primate: implications for conservation. Sci Rep 2015; 5: 14862.

61. Barelli C, Albanese D, Stumpf RM, Asangba A, Donati C, Rovero F, Hauffe HC. The gut microbiota communities of wild arboreal and ground-feeding tropical primates are affected differently by habitat disturbance. Msystems 2020; 5.

62. Trosvik P, Rueness EK, de Muinck EJ, Moges A, Mekonnen A. Ecological plasticity in the gastrointestinal microbiomes of Ethiopian Chlorocebus monkeys. Sci Rep 2018; 8: 1–10.

63. Kohl KD, Skopec MM, Dearing MD. Captivity results in disparate loss of gut microbial diversity in closely related hosts. Conserv Physiol 2014; 2: cou009.

64. Cameron A, Gould L. Fragment-Adaptive Behavioural Strategies and Intersite Variation in the Ring-Tailed Lemur (Lemur catta) in South-Central Madagascar. In: Marsh LK, Chapman CA (eds). Primates in Fragments SE - 16. Springer New York, 2013, pp 227–243.

65. Chi X, Gao H, Wu G, Qin W, Song P, Wang L, Chen J, Cai Z, Zhang T. Comparison of gut microbiota diversity between wild and captive bharals (Pseudois nayaur). BMC Vet Res 2019; 15: 1–8.

66. Hale VL, Tan CL, Niu K, Yang Y, Zhang Q, Knight R, Amato KR. Gut microbiota in wild and captive Guizhou snub-nosed monkeys, Rhinopithecus brelichi. Am J Primatol 2019; 81: e22989.

67. Clayton JB, Vangay P, Huang H, Ward T, Hillmann BM, Al-Ghalith GA, Travis DA, Long HT, Van Tuan B, Van Minh V. Captivity humanizes the primate microbiome. Proc Natl Acad Sci 2016; : 201521835.

68. Clayton JB, Al-Ghalith GA, Long HT, Van Tuan B, Cabana F, Huang H, Vangay P, Ward T, Van Minh V, Tam NA. Associations between nutrition, gut microbiome, and health in a novel nonhuman primate model. Sci Rep 2018; 8.

69. Nelson TM, Rogers TL, Carlini AR, Brown M V. Diet and phylogeny shape the gut microbiota of Antarctic seals: a comparison of wild and captive animals. Environ Microbiol 2013; 15: 1132–1145.

70. Greene L, Blanco MB, Rambeloson E, Graubics K, Fanelli B, Colwell RR, Drea CM. Gut microbiota of frugo-folivorous sifakas across environments. Anim Microbiome 2021.

71. Tsukayama P, Boolchandani M, Patel S, Pehrsson EC, Gibson MK, Chiou KL, Jolly CJ, Rogers J, Phillips-Conroy JE, Dantas G. Characterization of wild and captive baboon gut microbiota and their antibiotic resistomes. Msystems 2018; 3.

72. Narat V, Amato KR, Ranger N, Salmona M, Mercier-Delarue S, Rupp S, Ambata P, Njouom R, Simon F, Giles-Vernick T. A multi-disciplinary comparison of great ape gut microbiota in a central African forest and European zoo. Sci Rep 2020; 10: 1–15.

73. Xiao Y, Xiao G, Liu H, Zhao X, Sun C, Tan X, Sun K, Liu S, Feng J. Captivity causes taxonomic and functional convergence of gut microbial communities in bats. PeerJ 2019; 7: e6844.

74. Fujimura KE, Slusher NA, Cabana MD, Lynch S V. Role of the gut microbiota in defining human health. Expert Rev Anti Infect Ther 2010; 8: 435–454.

75. Clayton JB, Gomez A, Amato K, Knights D, Travis DA, Blekhman R, Knight R, Leigh S, Stumpf R, Wolf T. The gut microbiome of nonhuman primates: Lessons in ecology and evolution. Am J Primatol 2018; : e22867.

76. Cheng Y, Fox S, Pemberton D, Hogg C, Papenfuss AT, Belov K. The Tasmanian devil microbiome—implications for conservation and management. Microbiome 2015; 3: 1–11.

77. Borbón-García A, Reyes A, Vives-Flórez M, Caballero S. Captivity shapes the gut microbiota of Andean bears: insights into health surveillance. Front Microbiol 2017; 8: 1316.

78. Ma T, Villot C, Renaud D, Skidmore A, Chevaux E, Steele M. Linking perturbations to temporal changes in diversity, stability, and compositions of neonatal calf gut microbiota: prediction of diarrhea. ISME J 2020; : 1–13.

79. Chomel BB, Belotto A, Meslin F-X. Wildlife, exotic pets, and emerging zoonoses. Emerg Infect Dis 2007; 13: 6.

80. LaFleur M, Reuter KE, Hall MB, Rasoanaivo HH, McKernan S, Ranaivomanana P, Michel A, Rabodoarivelo MS, Iqbal Z, Rakotosamimanana N. Drug-Resistant Tuberculosis in Pet Ring-Tailed Lemur, Madagascar. Emerg Infect Dis 2021; 27: 977.

81. LaFleur M, Clarke TA, Reuter KE, Schaefer MS. Illegal Trade of Wild-Captured Lemur catta within Madagascar. Folia Primatol 2019; 90: 199–214.

82. Tong Q, Cui L-Y, Du X-P, Hu Z-F, Bie J, Xiao J-H, Wang H-B, Zhang J-T. Comparison of gut microbiota diversity and predicted functions between healthy and diseased captive Rana dybowskii. Front Microbiol 2020; 11: 2096.

83. Watson SE, Hauffe HC, Bull MJ, Atwood TC, McKinney MA, Pindo M, Perkins SE. Global change-driven use of onshore habitat impacts polar bear faecal microbiota. ISME J 2019; 13: 2916–2926.

84. Kaakoush NO. Insights into the role of Erysipelotrichaceae in the human host. Front Cell Infect Microbiol 2015; 5: 84.

85. Campbell TP, Sun X, Patel VH, Sanz C, Morgan D, Dantas G. The microbiome and resistome of chimpanzees, gorillas, and humans across host lifestyle and geography. ISME J 2020; 14: 1584–1599.

86. Nishida AH, Ochman H. A great-ape view of the gut microbiome. Nat Rev Genet 2019; 20: 195–206.

87. Greene LK, Williams C V, Junge RE, Mahefarisoa KL, Rajaonarivelo T, Rakotondrainibe H, O’Connell TM, Drea CM. A role for gut microbiota in host niche differentiation. ISME J 2020; : 1–13.

88. Sun Y, Sun Y, Shi Z, Liu Z, Zhao C, Lu T, Gao H, Zhu F, Chen R, Zhang J. Gut Microbiota of Wild and Captive Alpine Musk Deer (Moschus chrysogaster). Front Microbiol 2020; 10: 3156.

89. Liu J, Pu Y-Y, Xie Q, Wang J-K, Liu J-X. Pectin induces an in vitro rumen microbial population shift attributed to the pectinolytic Treponema group. Curr Microbiol 2015; 70: 67–74.

90. Liu J, Wang J-K, Zhu W, Pu Y-Y, Guan L-L, Liu J-X. Monitoring the rumen pectinolytic bacteria Treponema saccharophilum using real-time PCR. FEMS Microbiol Ecol 2014; 87: 576–585.

91. Dishman DL, Thomson DM, Karnovsky NJ. Does simple feeding enrichment raise activity levels of captive ring-tailed lemurs (Lemur catta)? Appl Anim Behav Sci 2009; 116: 88–95.

92. Mowry CB, Campbell JL, Mowry CB, Campbell JL. AZA Nutrition Advisory Group TAG/SSP Husbandry Notebook Nutrition Section Lemur catta (Ring-tailed lemur). 2001.

93. Zaneveld JR, McMinds R, Thurber RV. Stress and stability: applying the Anna Karenina principle to animal microbiomes. Nat Microbiol 2017; 2: 1–8.

94. Ahmed HI, Herrera M, Liew YJ, Aranda M. Long-term temperature stress in the coral model Aiptasia supports the “Anna Karenina principle” for bacterial microbiomes. Front Microbiol 2019; 10: 975.

95. Kohl KD, Dearing MD. Wild-caught rodents retain a majority of their natural gut microbiota upon entrance into captivity. Environ Microbiol Rep 2014; 6. doi:10.1111/1758-2229.12118.

96. Martínez-Mota R, Kohl KD, Orr TJ, Dearing MD. Natural diets promote retention of the native gut microbiota in captive rodents. ISME J 2020; 14: 67–78.

97. Grieneisen LE, Charpentier MJE, Alberts SC, Blekhman R, Bradburd G, Tung J, Archie EA. Genes, geology and germs: gut microbiota across a primate hybrid zone are explained by site soil properties, not host species. Proc R Soc B 2019; 286: 20190431.

98. Goodman SM, Benstead JP. Updated estimates of biotic diversity and endemism for Madagascar. Oryx 2005; 39: 73–77.

99. Ganzhorn JU, Lowry PP, Schatz GE, Sommer S. The biodiversity of Madagascar: one of the world’s hottest hotspots on its way out. Oryx 2001; 35: 346–348.

100. Dietrich M, Wilkinson DA, Soarimalala V, Goodman SM, Dellagi K, Tortosa P. Diversification of an emerging pathogen in a biodiversity hotspot: L eptospira in endemic small mammals of M adagascar. Mol Ecol 2014; 23: 2783–2796.

101. Jeffries CL, Tantely LM, Raharimalala FN, Hurn E, Boyer S, Walker T. Diverse novel resident Wolbachia strains in Culicine mosquitoes from Madagascar. Sci Rep 2018; 8: 1– 15.

102. Larsen PA, Hayes CE, Williams C V, Junge RE, Razafindramanana J, Mass V, Rakotondrainibe H, Yoder AD. Blood transcriptomes reveal novel parasitic zoonoses circulating in Madagascar’s lemurs. Biol Lett 2016; 12: 20150829.

103. Guiyoule A, Rasoamanana B, Buchrieser C, Michel P, Chanteau S, Carniel E. Recent emergence of new variants of Yersinia pestis in Madagascar. J Clin Microbiol 1997; 35: 2826–2833.

104. Bahram M, Hildebrand F, Forslund SK, Anderson JL, Soudzilovskaia NA, Bodegom PM, Bengtsson-Palme J, Anslan S, Coelho LP, Harend H. Structure and function of the global topsoil microbiome. Nature 2018; 560: 233–237.

105. Reed KE, Fleagle JG. Geographic and climatic control of primate diversity. Proc Natl Acad Sci 1995; 92: 7874–7876.

106. Mittermeier RA. Primate diversity and the tropical forest. Biodiversity 1988.

107. Melfi V. The appliance of science to zoo-housed primates. Appl Anim Behav Sci 2005; 90: 97–106.

108. Primates M, Altschul DM, Beran MJ, Bohn M, Call J, DeTroy S, Duguid SJ, Egelkamp CL, Fichtel C, Fischer J. Establishing an infrastructure for collaboration in primate cognition research. PLoS One 2019; 14: e0223675.

109. Johns T, Duquette M. Detoxification and mineral supplementation as functions of geophagy. Am J Clin Nutr 1991; 53: 448–456.

110. Krishnamani R, Mahaney WC. Geophagy among primates: adaptive significance and ecological consequences. Anim Behav 2000; 59: 899–915.

111. Won Y-J, Hallam SJ, O’Mullan GD, Pan IL, Buck KR, Vrijenhoek RC. Environmental acquisition of thiotrophic endosymbionts by deep-sea mussels of the genus Bathymodiolus. Appl Environ Microbiol 2003; 69: 6785–6792.

112. Tout J, Astudillo-García C, Taylor MW, Tyson GW, Stocker R, Ralph PJ, Seymour JR, Webster NS. Redefining the sponge-symbiont acquisition paradigm: sponge microbes exhibit chemotaxis towards host-derived compounds. Environ Microbiol Rep 2017; 9: 750–755.

113. Caravaggi A, Plowman A, Wright DJ, Bishop CM. The composition of captive ruffed lemur (Varecia spp.) diets in UK zoological collections, with reference to the problems of obesity and iron storage disease. J Zoo Aquarium Res 2018; 6: 41–49.

114. McPherson FJ. Normal blood parameters, common diseases and parasites affecting captive non-human primates. J Primatol 2013; 2: e112.

115. McKenney EA, Greene LK, Drea CM, Yoder AD. Down for the count: Cryptosporidium infection depletes the gut microbiome in Coquerel’s sifakas. Microb Ecol Health Dis 2017; 28: 1335165.

116. Schwitzer C, Mittermeier RA, Johnson SE, Donati G, Irwin M, Peacock H, Ratsimbazafy J, Razafindramanana J, Louis EE, Chikhi L. Averting lemur extinctions amid Madagascar’s political crisis. Science (80-) 2014; 343: 842–843.

117. Trevelline BK, Fontaine SS, Hartup BK, Kohl KD. Conservation biology needs a microbial renaissance: a call for the consideration of host-associated microbiota in wildlife management practices. Proc R Soc B 2019; 286: 20182448.

118. Tenhumberg B, Tyre AJ, Shea K, Possingham HP. Linking wild and captive populations to maximize species persistence: optimal translocation strategies. Conserv Biol 2004; 18: 1304–1314.

119. Mills JG, Weinstein P, Gellie NJC, Weyrich LS, Lowe AJ, Breed MF. Urban habitat restoration provides a human health benefit through microbiome rewilding: the Microbiome Rewilding Hypothesis. Restor Ecol 2017; 25: 866–872.

120. Song SJ, Amir A, Metcalf JL, Amato KR, Xu ZZ, Humphrey G, Knight R. Preservation methods differ in fecal microbiome stability, affecting suitability for field studies. MSystems 2016; 1.

121. Choo JM, Leong LE, Rogers GB. Sample storage conditions significantly influence faecal microbiome profiles. Sci Rep 2015; 5: 16350.

122. Caporaso JG, Lauber CL, Walters WA, Berg-Lyons D, Huntley J, Fierer N, Owens SM, Betley J, Fraser L, Bauer M. Ultra-high-throughput microbial community analysis on the Illumina HiSeq and MiSeq platforms. ISME J 2012; 6: 1621.

123. Bornbusch SL, Grebe NM, Lunn S, Southworth CA, Dimac-Stohl K, Drea C. Stable and transient structural variation in lemur vaginal, labial and axillary microbiomes: patterns by species, body site, ovarian hormones and forest access. FEMS Microbiol Ecol 2020; 96: fiaa090.

124. Quast C, Pruesse E, Yilmaz P, Gerken J, Schweer T, Yarza P, Peplies J, Glöckner FO. The SILVA ribosomal RNA gene database project: improved data processing and web-based tools. Nucleic Acids Res 2012; 41: D590–D596.

125. Yarza P, Yilmaz P, Pruesse E, Glöckner FO, Ludwig W, Schleifer K-H, Whitman WB, Euzéby J, Amann R, Rosselló-Móra R. Uniting the classification of cultured and uncultured bacteria and archaea using 16S rRNA gene sequences. Nat Rev Microbiol 2014; 12: 635.

126. Breiman L. Random forests. Mach Learn 2001; 45: 5–32.

127. Morton JT, Marotz C, Washburne A, Silverman J, Zaramela LS, Edlund A, Zengler K, Knight R. Establishing microbial composition measurement standards with reference frames. Nat Commun 2019; 10: 1–11.

128. Fedarko MW, Martino C, Morton JT, González A, Rahman G, Marotz CA, Minich JJ, Allen EE, Knight R. Visualizing’omic feature rankings and log-ratios using Qurro. NAR genomics Bioinforma 2020; 2: lqaa023.

129. Shenhav L, Thompson M, Joseph TA, Briscoe L, Furman O, Bogumil D, Mizrahi I, Pe’er I, Halperin E. FEAST: fast expectation-maximization for microbial source tracking. Nat Methods 2019; 16: 627.

