## Supplementary Material 1 for "Gut microbiota of ring-tailed lemurs (*Lemur catta*) vary across natural and captive populations and correlate with environmental microbiota" (from another version of this preprint: version 1, 2021-06-28)

### *1. Statistical results on alpha diversity in lemur gut microbiota*

The full statistical results for two metrics of alpha diversity (Shannon and Faith's Phylogenetic diversity) in lemur gut microbiota are shown in Tables S1-S4. For each metric, we (a) used a generalized linear mixed model (GLMM) to determine the predictive value of environment condition (wilderness; wild lemurs, captivity in Madagascar; $\text{captive}_{\text{M}}$ lemurs, and captivity in the U.S.; $\text{captive}_{\text{U.S.}}$ lemurs; Table S1 & S3) and setting (See Table 1 in the main text for names and descriptions of the 13 different settings; Table S2 & S4), and (b) Kruskal-Wallis rank sum tests and pairwise comparisons using Wilcoxon rank sum exact tests, with Benjamini-Hochberg adjustment for post-hoc and pairwise statistical comparisons.

**Table S1. Shannon diversity, by condition**

GLMM: Shannon diversity ~ condition + sex

| | Df | Deviance | AIC | F | p-value |
|---|---|---|---|---|---|
| <none> | | 29.243 | 192.27 | | |
| condition | 2 | 38.099 | 230.86 | 23.773 | 9.57E-10 |
| sex | 1 | 29.304 | 190.61 | 0.33 | 0.566 |

Kruskal-Wallis rank sum test

Kruskal-Wallis chi-squared = 30.694, df = 2, p-value = 2.162e-07

Pairwise comparisons using Wilcoxon rank sum exact test, with Benjamini-Hochberg adjustment:

| | captivity: Madagascar | captivity: US |
|---|---|---|
| captivity: US | 0.00023 | - |
| wilderness | 1.30E-07 | 0.635 |

**Table S2. Shannon diversity, by setting**

GLMM: Shannon diversity ~ setting + sex

| | Df | Deviance | AIC | F | p-value |
|---|---|---|---|---|---|
| <none> | | 21.291 | 155.18 | | |
| setting | 9 | 38.099 | 230.86 | 13.157 | 2.35E-15 |
| sex | 1 | 21.337 | 153.53 | 0.325 | 0.569 |

Kruskal-Wallis rank sum test

Kruskal-Wallis chi-squared = 89.371, df = 12, p-value = 6.535e-14

Pairwise comparisons using Wilcoxon rank sum exact test, with Benjamini-Hochberg adjustment:

| | AMO | BER | BEZ | DLC | FIH | ISO | IVO | LRC | NZP | NCZ | pet | RAN |
|---|---|---|---|---|---|---|---|---|---|---|---|---|
| BER | 0.777 | - | - | - | - | - | - | - | - | - | - | - |
| BEZ | 0.017 | 0.000 | - | - | - | - | - | - | - | - | - | - |
| DLC | 0.688 | 0.840 | 0.000 | - | - | - | - | - | - | - | - | - |
| FIH | 0.688 | 0.707 | 0.975 | 0.737 | - | - | - | - | - | - | - | - |
| ISO | 0.000 | 0.001 | 0.000 | 0.002 | 0.120 | - | - | - | - | - | - | - |
| IVO | 0.100 | 0.061 | 0.000 | 0.110 | 0.299 | 0.021 | - | - | - | - | - | - |
| LRC | 0.000 | 0.000 | 0.000 | 0.000 | 0.108 | 0.647 | 0.022 | - | - | - | - | - |
| NZP | 0.171 | 0.100 | 0.001 | 0.114 | 0.408 | 0.707 | 0.552 | 0.694 | - | - | - | - |
| NCZ | 0.863 | 0.778 | 0.463 | 0.647 | 0.843 | 0.071 | 0.197 | 0.061 | 0.361 | - | - | - |
| pet | 0.157 | 0.173 | 0.000 | 0.295 | 0.547 | 0.171 | 0.975 | 0.108 | 0.519 | 0.242 | - | - |
| RAN | 0.688 | 0.934 | 0.000 | 0.777 | 0.688 | 0.002 | 0.098 | 0.001 | 0.110 | 0.519 | 0.286 | - |
| TSI | 0.197 | 0.207 | 0.000 | 0.361 | 0.519 | 0.003 | 0.302 | 0.000 | 0.202 | 0.427 | 0.688 | 0.302 |

**Table S3: Faith's phylogenetic diversity, by condition**

GLMM: Faith's phylogenetic diversity ~ condition + sex

| | Df | Deviance | AIC | F | p-value |
|---|---|---|---|---|---|
| <none> | | 1361.5 | 810.62 | | |
| condition | 2 | 1438 | 815.43 | 4.415 | 0.013 |
| sex | 1 | 1361.6 | 808.64 | 0.018 | 0.891 |

Kruskal-Wallis rank sum test:

Kruskal-Wallis chi-squared = 10.383, df = 2, p-value = 0.005563

Pairwise comparisons using Wilcoxon rank sum exact test, with Benjamini-Hochberg adjustment:

| | captive$_{\text{M}}$ lemurs | captive$_{\text{U.S.}}$ lemurs |
|---|---|---|
| captive$_{\text{U.S.}}$ lemurs | 0.021 | - |
| wild lemurs | 0.022 | 0.056 |

**Table S4: Faith's phylogenetic diversity, by setting**

GLMM: Faith's phylogenetic diversity ~ setting + sex

| | Df | Deviance | AIC | F | p-value |
|---|---|---|---|---|---|
| <none> | 1075 | 786.59 | | | |
| setting | 9 | 1438 | 815.43 | 5.628 | 1.08E-06 |
| sex | 1 | 1075.3 | 784.63 | 0.041 | 0.839 |

Kruskal-Wallis rank sum test:

Kruskal-Wallis chi-squared = 64.518, df = 12, p-value = 3.347e-09

Pairwise comparisons using Wilcoxon rank sum exact test, with Benjamini-Hochberg adjustment:

| | AMO | BER | BEZ | DLC | FIH | ISO | IVO | LRC | NZP | NCZ | pet | RAN |
|---|---|---|---|---|---|---|---|---|---|---|---|---|
| BER | 0.019 | - | - | - | - | - | - | - | - | - | - | - |
| BEZ | 0.003 | 0.886 | - | - | - | - | - | - | - | - | - | - |
| DLC | 0.019 | 1.000 | 0.886 | - | - | - | - | - | - | - | - | - |
| FIH | 0.913 | 0.767 | 0.705 | 0.751 | - | - | - | - | - | - | - | - |
| ISO | 0.210 | 0.000 | 0.000 | 0.000 | 0.714 | - | - | - | - | - | - | - |
| IVO | 0.705 | 0.015 | 0.000 | 0.018 | 1.000 | 0.015 | - | - | - | - | - | - |
| LRC | 0.851 | 0.015 | 0.005 | 0.018 | 0.886 | 0.680 | 0.412 | - | - | - | - | - |
| NZP | 0.633 | 0.016 | 0.009 | 0.041 | 0.705 | 0.886 | 0.190 | 0.705 | - | - | - | - |
| NCZ | 0.061 | 0.310 | 0.251 | 0.251 | 0.600 | 0.012 | 0.018 | 0.106 | 0.120 | - | - | - |
| pet | 0.633 | 0.169 | 0.031 | 0.080 | 1.000 | 0.029 | 0.861 | 0.395 | 0.203 | 0.106 | - | - |
| RAN | 0.014 | 0.886 | 0.847 | 0.886 | 0.886 | 0.000 | 0.001 | 0.018 | 0.015 | 0.251 | 0.060 | - |
| TSI | 0.251 | 0.018 | 0.001 | 0.022 | 1.000 | 0.000 | 0.406 | 0.179 | 0.148 | 0.015 | 0.847 | 0.002 |

### *2. Statistical results on covariation between lemur gut and soil microbiota*

Table S4. Full statistical results (Wilcoxon rank sum exact tests, with Benjamini-Hochberg adjustment) for comparing mean proportion of soil-associated microbes in the gut microbiota of lemurs within and between the three conditions (wilderness, captivity in Madagascar, and captivity in the U.S.; Figure 9 in the main text).

| | captive$_{M}$ lemurs vs. captive$_{M}$ lemurs | captive$_{M}$ lemurs vs. captive$_{U.S.}$ lemurs | captive$_{M}$ lemurs vs. wild lemurs | captive$_{U.S.}$ lemurs vs. captive$_{M}$ lemurs | captive$_{U.S.}$ lemurs vs. captive$_{U.S.}$ lemurs | captive$_{U.S.}$ lemurs vs. wild lemurs | wild lemurs vs. captive$_{M}$ lemurs | wild lemurs vs. captive$_{U.S.}$ lemurs |
|---|---|---|---|---|---|---|---|---|
| captive$_{M}$ lemurs vs. captive$_{U.S.}$ lemurs | 8.90E-08 | - | - | - | - | - | - | - |
| captive$_{M}$ lemurs vs. wild lemurs | 2.00E-16 | 6.60E-13 | - | - | - | - | - | - |
| captive$_{U.S.}$ lemurs vs. captive$_{M}$ lemurs | 0.0367 | 0.00018 | 4.30E-11 | - | - | - | - | - |
| captive$_{U.S.}$ lemurs vs. captive$_{U.S.}$ lemurs | 2.20E-16 | 9.30E-05 | 0.136 | 1.60E-12 | - | - | - | - |
| captive$_{U.S.}$ lemurs vs. wild lemurs | 2.00E-16 | 2.00E-16 | 1.30E-05 | 4.80E-12 | 0.001 | - | - | - |
| wild lemurs vs. captive$_{M}$ lemurs | 0.8311 | 4.60E-15 | 2.00E-16 | 0.004 | 2.00E-16 | 2.00E-16 | - | - |
| wild lemurs vs. captive$_{U.S.}$ lemurs. | 2.00E-16 | 3.30E-10 | 0.084 | 3.30E-11 | 0.064 | 0.403 | 2.00E-16 | - |
| wild lemurs vs. wild lemurs | 3.50E-15 | 3.50E-15 | 2.00E-16 | 4.60E-10 | 0.00051 | 0.00011 | 2.00E-16 | 0.007 |

#### *3. Supplementary Figure 1: Differential abundance of soil microbes*

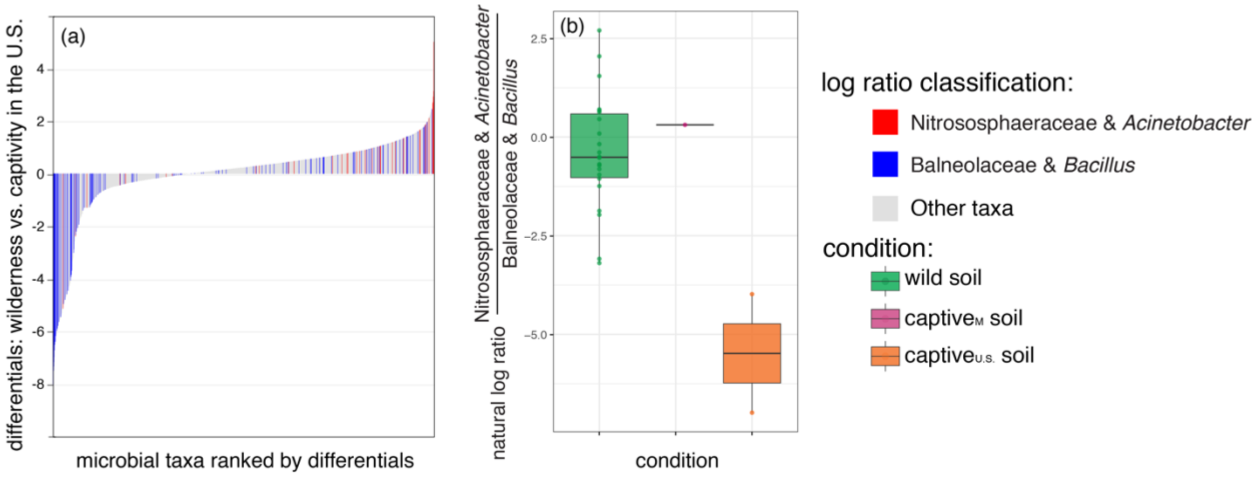


Figure S1. Differential abundance analysis of the family Nitrososphaeraceae and genus *Acinetorbacter* vs. the family Balneolaceae and genus *Bacillus* in soil microbiota. (a) Differential rank plot showing all microbial taxa (x-axis) ranked by their differentials (y-axis; the estimated log-fold changes for taxa abundances across sample groups) from wilderness vs. captivity in the U.S. Those taxa that are more abundant in the wild lemurs compared to captive lemurs in the U.S. appear on the right side of the plot whereas those that are less abundant in wild lemurs appear on the left side. Taxa belonging to family Nitrososphaeraceae and genus *Acinetorbacter* are highlighted in red whereas those in the family Balneolaceae and genus *Bacillus* are in blue (other soil taxa are in gray). (b) Natural log ratios of relative abundances of the two groups of taxa across soil from the three conditions.
